# The Impact of Serum on a Complex Synthetic Community Model of the Subgingival Microbiome

**DOI:** 10.1101/2025.10.07.681017

**Authors:** Lu Li, Matthew Smardz, Dam Soh, Philip D. Marsh, Anilei Hoare, Patricia I. Diaz

## Abstract

Despite rapid advances in characterizing the human microbiome, the ecological pressures shaping its transitions from healthy to diseased states remain poorly resolved. This is particularly true for periodontitis, a slow-progressing chronic inflammatory disease associated with well-defined shifts in the subgingival microbiome. Here, we report the development of a complex synthetic community model of the subgingival microbiome, designed for systematic interrogation of ecological factors that drive community restructuring. The model includes 22 prevalent and abundant subgingival species maintained in mucin-rich medium under microaerophilic, continuous culture conditions, in a chemostat. Using this system, we interrogated the impact of serum, as a surrogate for the inflammatory exudate, on community structure and function. Through integrated 16S rRNA gene sequencing, metatranscriptomics, and metabolomics, we found that serum was not required for a community with a periodontitis-like configuration to establish, but its presence intensified features of dysbiosis. Serum increased total biomass, promoted polymicrobial aggregate formation, promoted nitrogen and protein metabolism thereby modifying the environmental pH towards alkalinity, and introduced nitrosative stress. Serum also modified the community metatranscriptome in ways that paralleled microbiome activities in human periodontitis. Serum, however, decreased community diversity by disproportionally conferring a competitive advantage to the pathogen *Porphyromonas gingivalis*. This synthetic community model has revealed serum as a key nutritional pressure that modulates subgingival microbiome ecology and may perpetuate dysbiosis.

## Introduction

Recent omics-based studies of human associated microbiomes have advanced our understanding, in an unprecedented manner, of the disruptions in host-microbiome homeostasis that accompany a wide-spectrum of diseases [1–4]. In the oral cavity, the chronic inflammatory disease periodontitis is intricately linked to profound compositional and functional shifts in the microbiome community residing at the gingival crevice (the subgingival microbiome). Periodontitis-associated microbial communities are generally more diverse than those in health, enriched with numerous taxa—predominantly Gram-negative—and exhibit higher microbial biomass than health-associated communities [5–7]. The relationship between microbial dysbiosis and destructive inflammation of the periodontal tissues is thought to be part of a self-perpetuating cycle in which enrichment of pathobionts at the gingiva triggers a dysregulated inflammatory host response causing tissue inflammation and damage, which in turns alters the subgingival environment promoting further pathobiont enrichment [8, 9]. Understanding of the ecological drivers of subgingival community dysbiosis is, however, still incomplete, which has resulted in lack of successful therapies to restore communities to a health-compatible configuration [10].

The evolution of microbial communities during periodontitis is driven by interactions among its members and with the environment, ultimately generating a niche conducive to pathobiont expansion. Investigating these processes directly in humans remains difficult because periodontitis progresses slowly, and even with extensive longitudinal sampling it is not possible to disentangle the contributions of individual microorganisms or environmental factors to community dynamics. In previous studies, inoculation of natural oral or fecal microbiome communities into in vitro static systems enabled inquiries regarding the role of the nutritional environment on community composition [11–14]. While these models are valuable for probing factors that influence dysbiosis, natural communities vary across hosts and lack the manipulability required for mechanistic evaluation of the role of specific members or their genes as determinants of community outcomes. Synthetic communities, on the contrary, offer the advantage of being reproducible, allowing membership manipulations, and including well-characterized species. In previous work we employed synthetic communities of up to ten oral species to assess how the presence or absence of specific taxa influenced community composition[15, 16]. In addition, these studies coupled defined communities with a chemostat continuous culture system, in which nutrient influx rates can be modulated to support growth of slow-dividing species, thereby overcoming a known limitation of static batch models. This platform also enables repeated sampling for longitudinal tracking of communities and permits precise control of environmental parameters, allowing systematic evaluation of the influence of individual factors on community properties. Microbiome shifts associated with periodontitis development are well defined, at least from a taxonomic perspective [4, 6], with recent insight into the community transcriptional activities [17, 18]. Therefore, developing a complex synthetic community that models the subgingival microbiome represents a logical next step to rigorously dissect the ecological drivers of dysbiosis.

One of the main environmental influences thought to drive subgingival microbiome dysbiosis is gingival crevicular fluid (GCF), an exudate that bathes subgingival communities and contains serum-derived glycoproteins as well as tissue-produced inflammatory and breakdown products [19–21]. As inflammation ensues at the periodontal tissues, the flow rate of GCF increases [22, 23]. Support for the notion that the inflammatory exudate serves as a nutritional selective force that promotes dysbiotic shifts comes from previous studies with natural subgingival communities inoculated in closed static in vitro systems, in which the presence of serum, as a surrogate for GCF, promoted the enrichment of certain bacteria typical of inflamed periodontal pockets [11–13]. The manner in which serum modifies community metabolism and function, and whether this nutritional resource is the main ecological driver promoting the assembly of periodontitis-associated communities, are concepts still incompletely understood. Here, we describe the development of a complex synthetic subgingival community including 22 species maintained in a chemostat under continuous culture and demonstrate that, when coupled with omics analytical techniques, this model provides a powerful platform to dissect community interactions with specific environmental factors. We used this model to investigate the nutritional role of serum on community composition and function. Our work shows that serum, regardless of concentration, promoted communities with increased biomass and conferred a clear competitive advantage to the pathogen *Porphyromonas gingivalis*, with its enrichment leading to decreased community diversity. Elevated serum enhanced planktonic aggregate formation and switched community metabolism promoting ammonia production and a raise in the environmental pH, in part due to the activities of *P. gingivalis*. Higher serum also resulted in community-wide changes in expression of genes involved in protein and DNA repair, suggestive of a response to nitrosative stress, and modified community transcriptional activities in a manner that partly paralleled the microbiome functional activities in human periodontitis. Our studies suggest that serum acts as a nutritional pressure that modifies the subgingival microbiome recapitulating some of the features of dysbiotic communities in humans. Serum, however, does not support the highly diverse communities seen in many cases of periodontitis, suggesting that other ecological drivers also play a role in vivo, shaping communities into a periodontitis-like configuration.

## Methods

### Preparation of inoculum for synthetic community

Twenty-two bacterial species commonly found in the subgingival biofilm were selected based on their distribution in either periodontally healthy or periodontitis-affected individuals, as reported in our previous study [5]. The strains *Actinomyces oris* T14V, *Rothia dentocariosa* ATCC 17931, *Gemella haemolysans* ATCC 10379, *Streptococcus sanguinis* SK36, *Capnocytophaga gingivalis* ATCC 33624, and *Porphyromonas catoniae* F0037 were selected as periodontal health-associated species. *Corynebacterium matruchotii* ATCC 33806, *Veillonella parvula* PK1910, *Prevotella nigrescens* ATCC 33563, *Fusobacterium nucleatum subsp. nucleatum* ATCC 25586, *Eikenella corrodens* ATCC 23834, and *Lautropia mirabilis* ATCC 51599 were selected to represent core species. Finally, the periodontitis-associated species *Eubacterium brachy* ATCC 33089, *Filifactor alocis* ATCC 35896, *Streptococcus anginosus* ATCC 33397, and *Streptococcus constellatus* ATCC 27823, *Hoylesella oralis* ATCC 33269 (formerly *Prevotella oralis*), *Porphyromonas gingivalis* W83, *Porphyromonas endodontalis* ATCC 35406, *Prevotella melaninogenica* ATCC 25845, *Tannerella forsythia* ATCC 43037, and *Treponema denticola* ATCC 35405 were included (Fig. 1a). Each strain was maintained in the appropriate medium and cultured at 37°C under the appropriate atmospheric conditions (Supplementary Table 1).

**Fig. 1.**
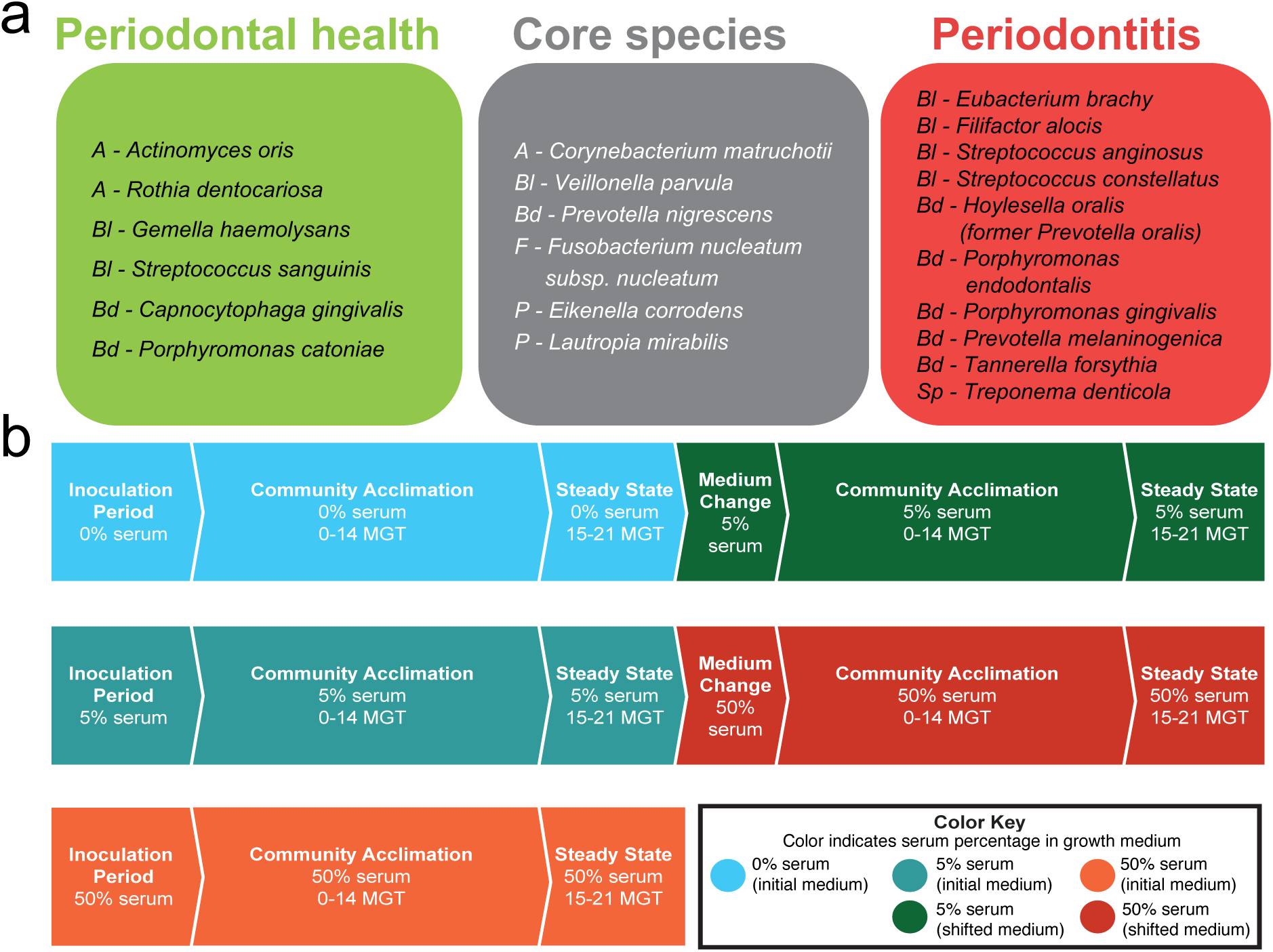
Overview of the synthetic community and experimental design for chemostat experiments. (a) Inocula for the synthetic community included twenty-two highly prevalent and abundant subgingival species associated with periodontal health, periodontitis, or non-associated (core species). Species are shown with a prefix denoting their phylum: *A* = *Actinomycetota*, *Bl* = *Bacillota*, *Bd* = *Bacteroidota*, *F* = *Fusobacteriota*, *P* = *Pseudomonadota*, *Sp* = *Spirochaetota*. (b) Flowchart for the three different experimental designs used to test the role of serum on community properties.

For chemostat community assembly, standardized frozen inocula were prepared from starter cultures grown in the specific medium required for each microorganism. Each starter culture was grown until reaching an optical density at 600 nm (OD_600_) of approximately 0.7, corresponding to mid exponential phase. Cultures were then normalized to OD_600_=0.7, equivalent to approximately 10^9^ cells mL^-1^. Aliquots of 20 mL were collected and centrifuged at 3,800 RPM for 15 min. The resulting pellets were resuspended in 500 μL of storage medium (growth medium supplemented with 10% glycerol). Bacterial suspensions were then transferred into cryovials, frozen at –80 °C for 24 h, and subsequently stored in liquid nitrogen until use.

### Continuous culture conditions

Continuous culture experiments were conducted in a Bioflow/CelliGen 115 Bioreactor (New Brunswick Scientific, Edison, NJ, USA) using frozen inocula of approximately 10^10^ cells, which were inoculated in 400 mL of mucin-serum [24]. The medium contained 2.5 mg mL^-1^ hog gastric mucin (Sigma-Aldrich, St. Louis, MO, USA), 2.5 mg mL^-1^ KCl, 2.0 mg mL^-1^ proteose peptone, 1.0 mg mL^-1^ yeast extract, 1.0 mg mL^-1^ trypticase peptone, 1.0 µg mL^-1^ cysteine·HCl, 5 µg mL^-1^ hemin, 10 µg mL^-1^ N-acetylmuramic acid (NAM), and was supplemented after autoclaving with heat-inactivated horse serum (Gibco, Thermo Fisher Scientific, Waltham, MA, USA) to final concentrations of 0%, 5%, or 50% (v/v). Antifoam 204 (Sigma-Aldrich, St. Louis, MO, USA) was added to a final concentration of 0.005% prior to autoclaving. Strain inoculation and community assembly were performed as previously described [16]. A microaerophilic atmosphere consisting of 2% O_2_, 5% CO_2_, 93% N_2_ was maintained throughout experiments. Temperature and pH were automatically controlled at 37 °C and 7.15 ± 0.15, respectively.

Cultures were monitored daily by measuring optical density at 600 nm (OD₆₀₀), dry weight, and by microscopic examination using differential contrast microscopy. In addition, redox potential (*E*h), dissolved oxygen (DO), pH, and the volume of acid and base added were recorded daily. Cultures were considered to have reached steady-state after 15 mean generation times (MGT), with evidence of sustained stability based on dry weights and *E*h measurements.

Two pilot experiments were performed to determine the optimal conditions for the growth of species with varying growth rates (Supplementary Fig. S1). Five-species communities, consisting of *P. gingivalis*, *F. nucleatum subsp. nucleatum*, *V. parvula*, *A. oris*, and *S. sanguinis* were established in continuous culture at two different dilution rates. The first protocol used a doubling time (Td) of 6.7 h, approximating that of *Streptococcus* spp. and *A. oris*, with a dilution rate (D) of 0.103 h⁻¹, and a flow rate (F) of 51.56 mL h^-1^. The second protocol was based on the experimental growth rate of *P. gingivalis* (Td = 15 h), using a D = 0.0462 h⁻¹, and a F = 23.21 mL h^-1^. Colony-forming unit (CFU) counts were obtained for *A. oris*, *S. sanguinis*, *F. nucleatum*, and *V. parvula*, while *P. gingivalis* abundance was assessed by quantitative PCR (qPCR) targeting 16S rRNA gene copies, four days after steady-state was achieved [16]. As shown in Supplementary Fig. S1, *P. gingivalis* exhibited low biomass under the 6.7 h Td protocol compared with the other four species. In contrast, under the 15 h Td condition, all species in the community reached cell densities above 10^7^ CFU mL^-1^. Based on these results, the slower growth rate protocol (Td = 15 h) was selected for subsequent experiments with the 22-species communities.

Three types of experiments were conducted to evaluate the effect of serum on the 22-species community (Fig. 1b). In the first, the community was established without serum and subsequently shifted to 5% serum after reaching theoretical steady-state (15-21 MGT), after which the culture was monitored until a second steady-state was achieved. In the second, the community was assembled in the presence of 5% serum and shifted to 50% serum after steady-state. In the third, the community was established and maintained at 50% serum. In all cases, samples were collected for community analysis: (i) for DNA analysis, 2 mL of culture were collected daily, centrifuged at 10,000 RPM for 10 minutes, and the pellets were stored at −80°C; (ii) for RNA analysis, 15 mL of culture obtained at steady-state was centrifuged at 4,200 RPM for 15 min, the pellets were suspended in 1,500 uL of RNA protect (Qiagen, Hilden, Germany), divided into three 500 uL aliquots, and stored at −80 °C; and (iii) for spent medium analysis, the supernatants from (ii) were filter-sterilized and stored at −80 °C until use.

The establishment of the community was shown to be reproducible, as demonstrated by repeating the experiment initiated with 5% serum and shifted to 50% serum in two independent runs. Community composition assessed by 16S rRNA gene sequencing showed consistent patterns across runs (Supplementary Fig. S2a, b). Day-to-day variation in species relative abundances was strongly correlated, with 17 of 22 species exhibiting Spearman correlation coefficients greater than 0.7 (Supplementary Fig. S2c). Principal coordinate analysis (PCoA) of Bray–Curtis beta-diversity further confirmed reproducibility, showing no significant differences between runs at the same serum concentration (PERMANOVA, P > 0.05; Supplementary Fig. S2d).

### Bacterial load determination

Genomic DNA was extracted from culture samples as previously described [6]. Briefly, samples were treated with lysozyme and proteinase K, followed by purification using a commercially available kit according to the instructions of the manufacturer (DNeasy Blood and Tissue kit; Qiagen, Hilden, Germany). DNA yield and purity were assessed with a NanoDrop spectrophotometer (Thermo Fisher Scientific, Waltham, MA, USA). Total bacterial load was quantified by real-time quantitative PCR (qPCR) using universal 16S rRNA primers and a TaqMan probe, as previously described [6]. Standard curves were generated to calculate the number of 16S rRNA gene copies per mL of culture at each time point.

### Evaluation of community composition via 16S rRNA gene sequencing

Community composition was evaluated by sequencing the V1-V2 regions of the 16S rRNA gene using primers 8F (5’-AGAGTTTGATCMTGGCTCAG-3’) and 361R (5’-CYIACTGCTGCCTCCCGTAG-3’) which incorporated Illumina MiSeq adapter sequences and single-end barcodes as previously described [25]. Amplicon libraries were pooled and sequenced using the MiSeq Reagent kit v3 (2 × 300 bp; Illumina, San Diego, CA, USA). Sequences are available at the Short Reads Archive (PRJNA1336263).

### 16S rRNA amplicon data processing and analysis

Raw sequences were processed using Mothur (v1.44.3) [26]. Paired-end reads were first merged into contigs, and contigs containing ambiguous bases, shorter than 150 bp, or with homopolymer stretches exceeding 10 bases were removed. Chimeric sequences were identified and filtered out using UCHIME [27]. After quality control, a total of 7,596,297 sequences were retained (mean ± SD per sample 75,211 ± 48,252) for downstream analysis. Taxonomic classification was performed in Mothur using the k-nearest neighbor (knn) algorithm with k = 1, based on BLAST alignment to a curated reference set from the Human Oral Microbiome Database (v15.1) [28] comprising the 22 species included in the synthetic community. Reads not mapping to the reference sequences were excluded. Microbial composition was assessed using relative abundances. Finally, 7,596,297 sequences were retained (mean ± SD per sample: 75,211 ± 48,252) for downstream analyses.

Alpha diversity was quantified using the Shannon index to assess community evenness. Beta diversity was evaluated based on Bray–Curtis distances calculated on species relative abundances and visualized using principal coordinate analysis (PCoA). To identify species most strongly associated with the compositional shifts induced by serum, Spearman’s correlations were calculated between the relative abundance of each species and the first two principal coordinates (PCo1 and PCo2). Species showing significant correlations to PCo1 were highlighted as key contributors to the observed shifts. Differential abundances of species across steady-states were assessed using the Kruskal–Wallis test followed by Bonferroni post hoc correction.

### Evaluation of community transcriptional activity via metatranscriptomics

RNA was isolated from pellets resuspended in RNA protect (Qiagen) using the RNeasy Mini Kit (Qiagen, cat. no. 79254) following the manufacturer’s protocol with two modifications. First, pellets were resuspended in RLT buffer with β-mercaptoethanol and transferred to silica bead tubes (Lysing Matrix B, 2 mL; MP Biomedicals, Irvine, CA, USA; cat. no. 116911100), followed by lysis in a FastPrep-24 instrument (MP Biomedicals) for 20 s. Second, after the first RW1 wash step, on-column DNase digestion was performed using the RNase-Free DNase Set (Qiagen, cat. no. 79256; 10 μL DNase I in 70 μL Buffer RDD) for 15 min at room temperature. The protocol was then resumed, and RNA was eluted in nuclease-free, molecular biology-grade water.

RNA was stored at –80 °C until sequencing. RNA integrity was assessed with an Agilent Fragment Analyzer, and concentrations were determined using a Qubit fluorometer (Thermo Fisher Scientific). For library preparation, 100 ng of total RNA per sample was processed with the Illumina RiboZero Total Stranded RNA Library Prep Kit, including ribosomal RNA removal, according to the manufacturer’s protocol. Final libraries were pooled to a concentration of 10 nM determined by the QuantaBio Universal qPCR kit. After final dilution and denaturing, the pooled library was loaded in one lane at a concentration of 175 pM for 50 base pair paired-end sequencing in an Illumnia NovaSeq 6000 platform at the University at Buffalo Genomics and Bioinformatics Core (Buffalo, NY, USA). Sequences are available at the Short Reads Archive (PRJNA1336263).

### Metatranscriptomic data processing and analysis

Raw RNA sequencing reads were processed with Kneaddata (0.12.2) [29] for quality trimming, filtering, and removal of potential contaminant sequences by mapping against the human reference genome (GRCh37/hg19) [30]. After filtering, reads were aligned using Bowtie2 (2.5.4) [31] to a custom database comprising the reference genomes of the 22 species included in the synthetic community, which were retrieved from the Human Oral Microbiome Database (HOMD, v15.1). Gene counts were extracted from the alignment files with Samtools (v1.21) [32] based on annotated coding sequences and normalized to reads per kilobase (RPK) to account for gene length. For functional interpretation across species, genes were assigned to UniRef90 gene families using DIAMOND (2.0.15) [33] and subsequently aggregated into KEGG orthologies (KOs), KEGG pathways [34], and Gene Ontology (GO) terms [35].

Metatranscriptomic profiles were normalized to relative abundance, calculated as the proportion of RPK assigned to a given feature relative to the total RPK per sample. To evaluate shifts in the composition of transcriptionally active species at steady-state, Bray–Curtis beta diversity was assessed on transcripts grouped by species and visualized with PCoA. Correlation analysis with the principal coordinates was applied following the same approach used for 16S data, to identify species most strongly associated with serum-induced shifts. Alterations in the total transcriptional activity of individual species across steady-states were further evaluated using the Kruskal–Wallis test with Bonferroni post hoc correction.

Community-wide functional analysis was performed on aggregated functional profiles (e.g. UniRef90 gene families, KOs, KEGG pathways, and GO terms) to characterize the overall functional activity of the community. Beta diversity of these functional profiles was assessed using Bray–Curtis distances and visualized with PCoA, while differential expression was identified with DESeq2 [36]. Redundant GO terms were reduced with REVIGO [37].

Since community-level transcriptomic profiles reflect both variation in the abundance of metabolically active species and changes in gene expression within individual species, a complementary species-specific analysis was also performed. In this analysis, the RPK value of each gene was normalized to the total transcriptional output of its corresponding species, thereby controlling for differences in species abundance. This approach enabled the detection of genuine regulatory responses within taxa rather than changes attributable solely to shifts in community composition. Differential expression of genes within each species was identified using DESeq2, and differentially expressed biological processes (BPs) were likewise determined for the six species that showed the strongest transcriptional responses to serum, each exhibiting more than 20% of their expressed genes differentially regulated at higher serum concentrations.

### Metabolomics of spent media

Spent medium samples collected and stored as described above were used for metabolomic analysis. Metabolomic profiling was performed at the Southeast Center for Integrated Metabolomics (SECIM), University of Florida Health (Gainesville, FL, USA). Samples underwent protein precipitation, followed by collection and drying of the supernatant. Dried extracts were reconstituted for analysis in both positive and negative ionization modes, each run as separate injections. Global metabolomics profiling was carried out on a Thermo Q-Exactive Orbitrap mass spectrometer equipped with a Dionex UHPLC and autosampler. All samples were analyzed under heated electrospray ionization (positive and negative) at a mass resolution of 35,000 (m/z 200). Separation was achieved on an ACE 18-pfp column (100 × 2.1 mm, 2 µm) with mobile phase A (0.1% formic acid in water) and mobile phase B (acetonitrile), a flow rate of 350 µL/min, and a column temperature of 25 °C. Injection volumes were 4 µL for negative ions and 2 µL for positive ions. Each batch included extraction blanks, quality controls, and samples. Internal and injection standards were added to all samples to monitor batch reproducibility.

Data from positive and negative ionization modes were analyzed separately. A total of 1,663 features were detected in positive mode and 2,323 in negative mode. Feature detection, deisotoping, alignment, and gap filling were performed in MZmine [38]. Adducts and complexes were identified and removed. Features were annotated by searching against the SECIM’s internal retention time library of approximately 1,100 standards. Features lacking confident annotation were further queried against the Human Metabolome Database [39] to assign putative identities based on m/z, ionization mode, and molecular weight tolerance (±5 ppm).

Processed data were imported into MetaboAnalyst 5.0 [40] for statistical analysis. Features with >80% missing values were removed, and remaining missing values were imputed with the feature mean. Data were filtered by relative standard deviation (RSD = SD/mean), normalized by sum, log-transformed, and autoscaled using the default settings. Differential features across serum conditions in both ionization modes were identified within MetaboAnalyst. Based on the normalized metabolite abundances, community-level metabolic differences among the three serum conditions (0%, 5%, and 50%) were evaluated by principal component analysis (PCA) using Euclidean distance. Statistical significance was assessed using PERMANOVA followed by Bonferroni post hoc correction for multiple comparisons.

### Measurement of protein concentration of fresh media

Fresh hog mucin medium was prepared as described above with 0%, 5%, and 50% (v/v) heat-inactivated horse serum added after autoclaving. Protein concentrations in the media were quantified using the Pierce BCA Protein Assay Kit (Thermo Fisher Scientific) and measured with a Genesys 30 spectrophotometer (Thermo Fisher Scientific) using the manufacturer-supplied cuvettes.

### Determination of ammonium/ammonia concentrations in spent media

The concentration of ammonium/ammonia in spent media at steady-state was quantified using a colorimetric assay kit (Abcam, Cambridge, UK). Samples were thawed from −80°C storage and diluted as required to ensure their values fell within the linear range of the standard curve. A standard curve was generated using an ammonium chloride standard solution and sample concentrations were determined by interpolation from the standard curve.

### Growth of *F. nucleatum* and *P. gingivalis* as monocultures under different serum concentrations

To evaluate the effect of serum on *F. nucleatum* and *P. gingivalis*, both species were cultured in mucin medium, prepared as described above, supplemented with 0%, 5%, or 50% (v/v) heat-inactivated horse serum. The pH of the medium was adjusted to 7.0 before inoculation. Each species was inoculated at a density of 10⁷ cells mL⁻¹ and incubated anaerobically at 37 °C for seven days. Cultures were sampled daily inside the anaerobic chamber. Biomass of each species was quantified via qPCR, and the pH of culture media was measured using a pH meter (Accumet AB15, Fisherbrand, Hampton, NH, USA).

## Results

### Serum promotes greater biomass and inter-microbial assemblage formation

We developed a defined-inoculum polymicrobial community model in continuous culture consisting of 22 prevalent species of subgingival plaque (Fig. 1a), selected based on previous human studies [41]. The synthetic community was established under controlled conditions, including a consistent pH of 7.15± 0.15, a microaerophilic atmosphere (2% O_2_), and using a mucin-based basal medium [24], to replicate the oral environment, as nutritional source. Heat-inactivated serum, to a final concentration of 5% or 50% by volume, was introduced during initial community assembly or at steady-state to assess the impact of this nutritional substrate on the community properties (Fig. 1b).

Given that gingival inflammation and exposure of subgingival plaque to a higher flow rate of GCF is associated with increased microbiome biomass [6], we first evaluated the effect of serum on total bacterial load (Fig. 2a). We found that serum, regardless of concentration, significantly increased, by about one log, the total biomass compared to serum-free conditions. We next performed microscopic examination of fresh planktonic culture samples to determine whether serum influenced the physical organization and appearance of microbial cells. Microscopic examination revealed cultures consisted of microbial aggregates comprised of cells with diverse morphologies (Fig. 2b). These planktonic aggregates occurred in the absence and presence of serum and were reminiscent of the multi-species clusters that commonly occur in the planktonic salivary phase within the oral cavity [42]. Notably, although total load was similar between 5% and 50% serum (Fig. 2a), large dense aggregates characterized communities in the highest serum concentration (Fig. 2b), suggesting that an excess of serum components enhances the physical interactions among subgingival bacteria promoting the formation of structured microbial assemblages.

**Fig. 2.**
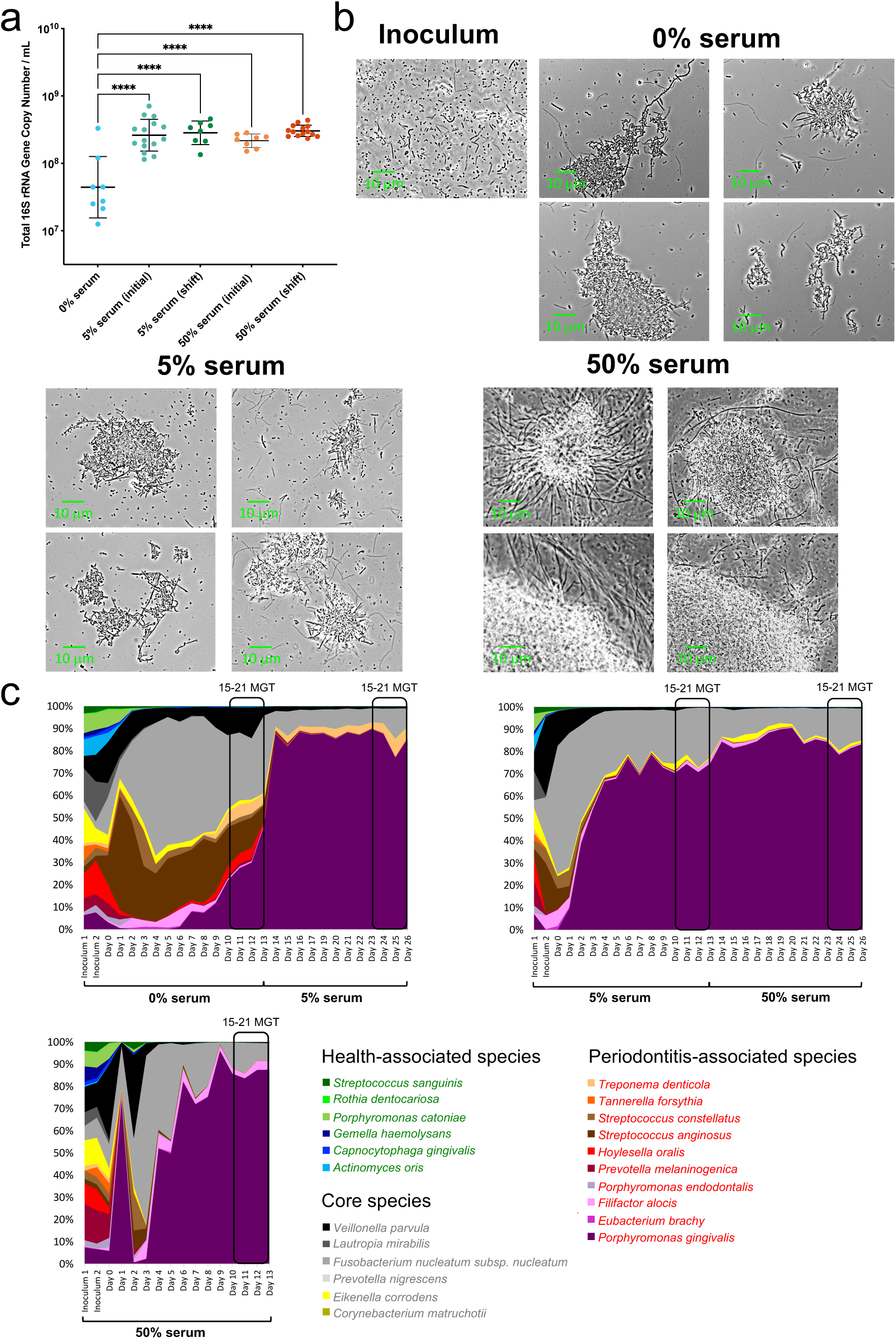
Effect of serum on community biomass, microbial assemblages and species abundances. (a) Total bacteria load at each steady state quantified using qPCR. Significance of differences between groups was determined using one-way ANOVA, with Bonferroni post hoc analysis (\*\*\*\**P* < 0.0001). (b) Phase-contrast micrographs of steady-state cultures showing microbial aggregate formation under different serum concentrations. (c) Community taxonomic composition during the assembly period and theoretical steady state phases (15-21 mean generation times, MGT) as characterized via 16S rRNA sequencing. For the condition initiated with 5% serum and shifted to 50% serum, the plot represents the average of two independent runs; the results of individual runs are shown in Supplementary Fig. S2.

### Serum affects polymicrobial community structure

To monitor daily community composition, we employed 16S rRNA gene sequencing (Supplementary Table S2). As shown in Fig. 2c and Supplementary Fig. S3, species distributions in the inoculum were relatively even. However, when growing in the basal mucin medium (no serum), community evenness dropped during the first 4 days in culture, with 8 species with a relative abundance greater than 1% dominating the community for the following days, until steady-state. These species were core species or periodontitis-associated taxa, while health-associated commensals were present but in low abundance (Supplementary Table S2). Introduction of serum, at any concentration, and either during initial community assembly or at steady-state, affected community structure further decreasing evenness and promoting the dominance of the periodontitis-associated species *Porphyromonas gingivalis*, only accompanied at greater than 1% abundance by *Fusobacterium nucleatum subsp. nucleatum*, in addition to *Filifactor alocis* under 50% serum.

A comparison of steady-state community structure via Bray-Curtis distances revealed distinct clustering between serum-free and serum-supplemented conditions (Fig. 3a, PERMANOVA, *P*=0.002 for both 0% vs. 5% serum and 0% vs. 50% serum). The timing of serum addition, whether during assembly or at steady-state, did not have a significant effect on community structure. However, communities grown at different serum concentrations (5% vs. 50%) showed statistically significant differences in composition (PERMANOVA, *P*=0.02) although clusters overlapped suggesting a lower effect size than when comparing to the no-serum condition (Fig. 3a). An analysis of the species driving these shifts showed that *P. gingivalis* was the main beneficiary of having serum in the growth medium, at the expense of depletion of certain core and periodontitis-associated species (Fig. 3a). Additionally, analysis of changes in proportions of individual species showed that *P. gingivalis* was the only species enriched in 5% serum when compared to no serum, with its levels remaining the same in 50% serum as in 5%. A higher serum concentration, however, supported higher proportions of the periodontitis-associated species *Filifactor alocis* and the health-associated *Gemella haemolysans*. Most other species were depleted under serum (Fig. 3b).

**Fig 3.**
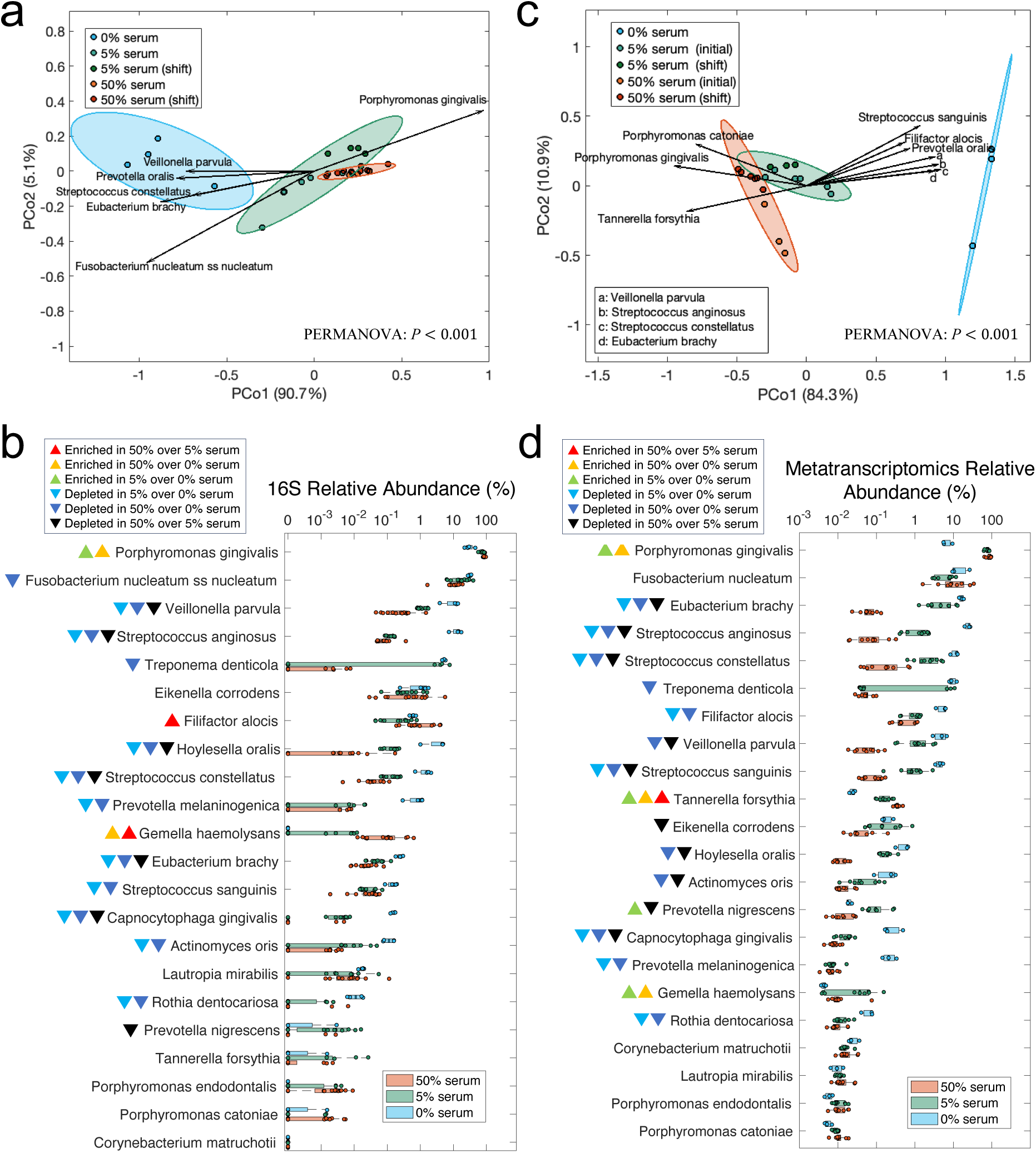
Effect of serum on steady-state microbial community composition, as analyzed through 16S rRNA sequencing and metatranscriptomics. (a) Principal coordinate analysis (PCoA) of Bray-Curtis beta-diversity based on 16S rRNA data across different serum concentrations at steady states. Group-level differences were assessed with PERMANOVA test with Bonferroni correction as post hoc analysis. P-values: 0.063 for 5% serum (initial) vs. 5% serum (shift); 1 for 50% serum (initial) vs. 50% serum (shift); 0.002 for 0% serum vs. 5% serum; 0.002 for 0% serum vs. 50% serum; 0.02 for 5% serum vs. 50% serum. Arrows indicate species significantly correlated with PCo1 (*P*<0.05), with arrow direction and length reflecting Spearman’s correlation between species relative abundances and the PCoA axes. (b) Box plots of 16S rRNA relative abundance changes across serum concentrations. Significance of differences was determined using the Kruskal–Wallis test with Bonferroni correction for multiple comparisons; colored triangles denote enriched species. (c) PCoA of Bray-Curtis beta-diversity based on species metatranscriptomics relative abundances across serum concentrations. PERMANOVA p-values: 0.334 for 5% serum (initial) vs. 5% serum (shift); 0.112 for 50% serum (initial) vs. 50% serum (shift); 0.014 for 0% serum vs. 5% serum; 0.017 for 0% serum vs. 50% serum; 0.005 for 5% serum vs. 50% serum. Arrows represent species significantly correlated with PCo1 (*P*<0.05) by the Spearman’s correlation. (d) Box plots of metatranscriptomic relative abundance changes across serum concentrations. Statistical significance was determined using the Kruskal–Wallis test with Bonferroni correction; colored triangles denote enriched species.

Since 16S rRNA gene sequencing does not distinguish between metabolically active and inactive species, we performed metatranscriptomic sequencing to evaluate transcriptionally active species at steady-state. This analysis confirmed the global-scale compositional shifts observed via 16S rRNA gene sequencing with *P. gingivalis* as the main beneficiary of serum introduction (Fig. 3c). Fig. 3d shows levels for transcriptionally active individual species across conditions showing that all 22 species were detected at steady-state, demonstrating that the model successfully sustains a diverse and functionally active microbial community. Interestingly, while *T. forsythia*, a species strongly associated with periodontitis, exhibited low abundance by 16S rRNA gene sequencing, it was moderately abundant in the metatranscriptomics data, with its transcriptional activity increasing proportionally to serum concentration. Other species showed comparable trends between the metatranscriptomics and 16S data, with most species decreasing in abundance in the presence of serum. *F. nucleatum*, an abundant component of dental plaque in both health and periodontitis did not change its relative abundance upon serum addition according to RNA sequencing, consistent with its role as a core subgingival species, able to thrive under both periodontal health and disease environments. Overall, these taxonomic analyses show that the community growing without serum was more diverse and dominated by core species and periodontitis-associated taxa, while introduction of serum as a nutrient source changed diversity and community structure promoting the competitive dominance of *P. gingivalis* with minor enrichment of *F. alocis* and *T. forsythia*, as shown by 16S and RNASeq analysis, respectively.

### Serum promoted community-wide metabolic shifts increasing alkaline and decreasing acidic end-products

We observed a notable increase in acid consumption in the chemostat system under serum, especially at 50% (Fig. 4a). Given that the basal pH of different media did not differ and that the system was designed to maintain a pH of 7.15±0.15, this increase in acid utilization suggested a metabolic shift towards production of less acidic and more alkaline fermentation end-products. To investigate this, we performed LC-MS analysis of spent media from steady-state cultures. Analysis of global metabolomic profiles showed the no serum, 5%, and 50% serum communities as distinct (Fig. 4b). Volcano plots showed a small number of metabolites differentially abundant in the comparison of spent media from no serum versus 5% and a larger number in the 5 to 50% serum comparison (Fig. 4c). This analysis showed that strong acidic end-products such as lactate, 2-furoic acid, mesylate, and oxalic acid were depleted in conditions with serum, whereas basic and amine-containing metabolites, including dimethylethanolamine, N-acetylputrescine, and L-lysine were enriched. Detailed differential metabolites are provided in Supplementary Table S3. Since the LC-MS technique used did not allow direct measurement of ammonia, a key alkaline by-product of amino acid metabolism, we next employed a colorimetric assay to quantify ammonia concentrations in the spent media. This analysis showed that ammonia levels were significantly elevated in steady-state communities grown with serum (5% and 50%) compared to serum-free conditions, with a trend towards higher levels at 50% serum (Fig. 4d). In summary, these results suggest serum induces metabolic shifts increasing ammonia production and depleting acidic end-product formation by the microbial community.

**Fig 4.**
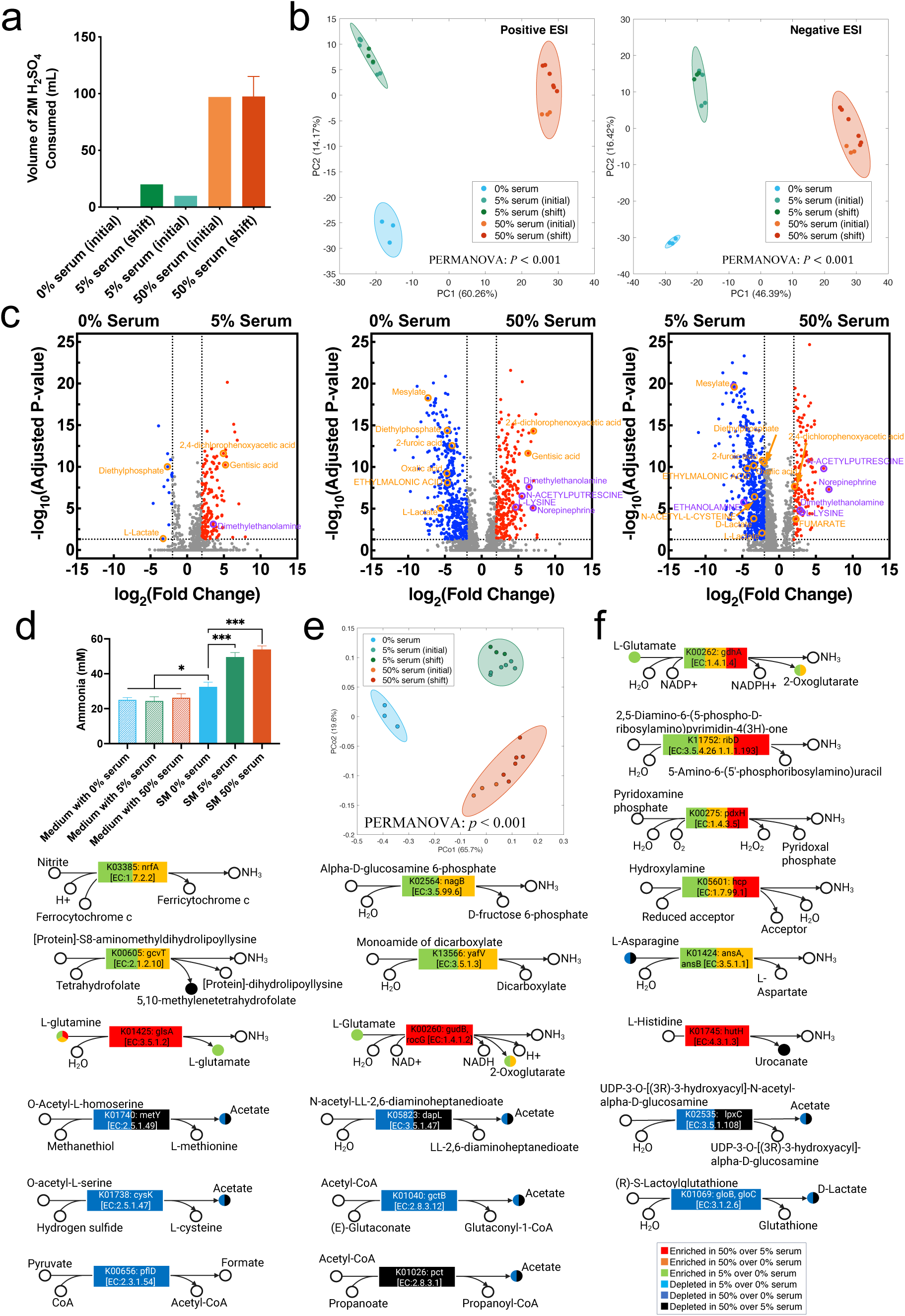
Effect of serum concentration on community metabolism, end-products and environmental pH. (a) Acid consumption at steady state in the chemostat system for each serum concentration. (b) Principal component analysis (PCA) of metabolomic peaks (positive and negative ESI modes) across steady-state samples with different serum concentrations. PERMANOVA tests were used to evaluate group-level differences, with Bonferroni correction. P-values: 0.005 for 0% vs. 5% serum; 0.011 for 0% vs. 50% serum; *P*<0.001 for 5% vs. 50% serum in positive ESI mode; and 0.013 for 0% vs. 5% serum; 0.011 for 0% vs. 50% serum; *P*<0.001 for 5% vs. 50% serum in negative ESI mode. (c) Volcano plots of metabolites detected by LC–MS showing differential abundance across serum concentrations. Red dots indicate metabolites enriched in higher serum, while blue dots indicate metabolites depleted. Acidic products (pKa<4) are highlighted in orange and basic products (pKa>9) in purple. Metabolites annotated using the SECIM internal reference standards database are shown in uppercase, whereas those queried against the Human Metabolome Database (HMDB) are shown in lowercase. Group-level differences were assessed using PERMANOVA with Bonferroni correction. (d) Bar plot of ammonia concentrations measured by a colorimetric assay in uninoculated medium and spent media (SM) at steady state across different serum concentrations. Significance of differences between conditions was evaluated using one-way ANOVA with Bonferroni correction (\**P*<0.05, \*\*\**P*<0.001). (e) Principal coordinate analysis (PCoA) of metatranscriptomic KEGG orthologies (KOs) based on Bray-Curtis distances for steady-state samples. Group-level differences were assessed using PERMANOVA with Bonferroni correction. P-values: 0.006 for 0% vs. 5% serum, 0.007 for 0% vs. 50% serum, and P < 0.001 for 5% vs. 50% serum. (f) KOs involved in ammonia production (upregulated by serum) and acidic product formation (downregulated by serum) identified by DESeq2. Colored circles indicate the corresponding metabolites detected as differentially abundant by LC–MS (panel c).

The shifts in metabolite profiles with serum suggested a potential change in transcriptomic activity related to metabolism. To explore this further, community-wide metatranscriptomes were annotated using KEGG orthologies (KOs). Principal Coordinate Analysis revealed distinct clustering of communities growing in no serum, 5% and 50%, confirming serum altered the community metabolic activities in a concentration dependent manner (Fig. 4e). Among KOs upregulated by serum were several genes involved in reactions that result in production of ammonia, while KOs associated with acidic end-product formation were downregulated (Fig. 4f). Notably, genes critical to nitrite reduction, such as K03385 (*nrfA*, nitric oxide reducatse), and nitrosative stress protection, including K05601 (*hcp*, hybrid cluster protein), were significantly upregulated in by serum, as were genes involved in glutamine and histidine catabolism, all contributing to ammonia production. In contrast, genes responsible for acetate and lactate production were downregulated in response to serum, with metabolomics confirming depletion of these end-products in spent media (Fig. 4f). Detailed differential KOs are provided in Supplementary Table S4. Furthermore, an analysis of KEGG pathways showed that serum was associated with enrichment of nitrogen metabolism, partly through upregulation of amino acid degradation pathways, including histidine metabolism (Supplementary Fig. S4). Since these results suggested increase protein catabolism under serum, we next measured the total protein content of the uninoculated media used in the 3 conditions tested, which showed 5% and 50% serum had ∼2-fold and ∼12-fold more protein, respectively, than the no serum medium (Fig. 5a). Serum was also associated with an increase in glycosaminoglycan and other glycan degradation suggesting the community could utilize these components of serum (Supplementary Fig. S4). Collectively, these findings show serum induces a metabolic shift favoring protein and nitrogen metabolism and ammonia production, while suppressing acidic end-product formation.

**Fig 5.**
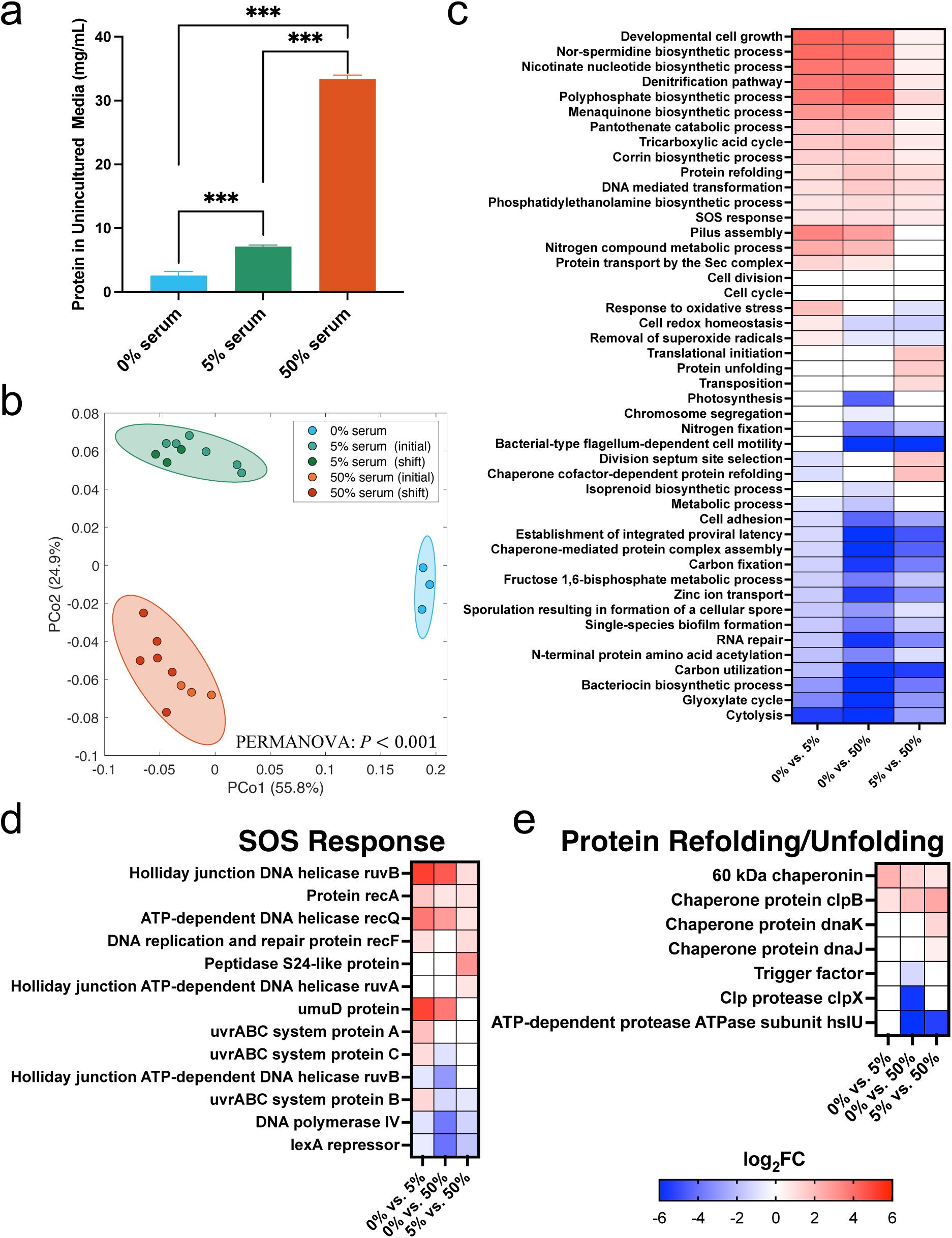
Effect of serum concentration on community-level transcriptomic activities. (a) Bar plot showing the total protein content of uninoculated media with varying serum concentrations. Significance of differences was evaluated using one-way ANOVA with Bonferroni correction (\*\*\**P*<0.001). (b) Principal coordinate analysis (PCoA) of Bray–Curtis beta-diversity based on metatranscriptomic data grouped by Gene Ontology (GO) terms across different serum concentrations. Group-level differences were assessed using PERMANOVA test with Bonferroni correction. P-values: 0.017 for 0% vs. 5% serum, 0.005 for 0% vs. 50% serum, and 0.001 for 5% vs. 50% serum. (c) Heatmap of log2 fold-changes (log2FC) of differentially expressed biological processes (BPs) across serum concentrations. (d–e) Differentially expressed UniRef90 gene families involved in (d) SOS response, and (e) protein refolding/unfolding. Differential expression across panels (c–e) was determined using DESeq2 (log2FC >0.3785 or <–0.3785, FDR<0.05).

### Serum modulates transcriptomic activities inducing a stress response

To gain broader insights into functional changes beyond metabolic functions, we annotated metatranscriptomic data using Gene Ontology (GO) terms. Principal Coordinate Analysis (PCoA) of the metatranscriptomic GO profiles showed significant differences between the no-serum, 5% and 50% serum conditions, again confirming serum influenced community-wide gene expression in a concentration dependent manner (Fig. 5b). Differential expression analysis identified several biological processes (BPs) that varied across serum concentrations (Fig. 5c). Notably, serum promoted the upregulation of the SOS response and protein refolding/unfolding pathways, indicating the community was responding to a stress. Genes involved in DNA repair, such as *recA*, *recQ*, *recF*, *ruvA*, *ruvB*, and the *uvrABC* system, were significantly upregulated, while the critical SOS repressor *lexA* was downregulated, pointing to a response to DNA damage (Fig. 5d). Additionally, several chaperones involved in protein maintenance, such as *clpB* and *dnaK*, were enriched in serum, reflecting increased proteostasis activity in response to stress (Fig. 5e).

Interestingly, the denitrification pathway was upregulated in response to serum (Fig. 5c). Serum is known to have nitrate and nitrite at concentrations in the μM range [43, 44], and higher levels of *nrfA*, a key enzyme in nitrite ammonification, were shown to be induced in the community by serum, suggesting an increase in nitrite respiration (Fig. 4f). Production of nitric oxide and reactive nitrogen species (RNS) during nitrite respiration could be, however, a source of DNA and protein damage, potentially explaining the observed upregulation in protective mechanisms, such as DNA repair and protein refolding. When considering other possible sources of DNA and protein damage, we observed that genes directly involved in oxidative stress were inconsistently upregulated under 5% serum and actually downregulated under 50% serum (Supplementary Fig. S5). The downregulation of oxidative stress response aligns with a very reduced redox potential (*Eh*) in cultures under all conditions with serum (Supplementary Fig. S6), despite the constant incoming flow of oxygen into the chemostat. Therefore, nitrosative stress was likely the source of stress under serum.

Another GO biological process upregulated by 50% serum was transposition (Fig. 5c), driven by the enrichment of several transposase and recombinase genes (Supplementary Table 5). This may be connected to the stress response, as higher ability to mobilize transposable elements could be advantageous conferring plasticity to species in the community. Other GO biological processes upregulated included cell growth, cell cycle, cell division, translation initiation (Fig. 5c), consistent with the higher biomass of communities under serum. Corrin biosynthesis, which forms the core for corrinoid cofactors such as cobalamin, was also upregulated under serum (Fig. 5c).

### Serum induces species-specific changes in transcriptomic activity

We further investigated species-specific transcriptomic responses to serum by normalizing gene expression within each species to its total transcriptomic output. This allowed us to identify differentially regulated genes within each species, independent of changes in community composition across conditions. Fig. 6a shows the percentage of differentially expressed genes within each species. Notably, six species, the core species *F. nucleatum*, the periodontitis-associated *P. gingivalis*, *Streptococcus constellatus*, *Eubacterium brachy*, and *F. alocis*, and the health-associated commensal *Streptococcus sanguinis*, exhibited the most substantial transcriptomic shifts, with over 20% of their expressed genes differentially regulated in response to higher serum concentration. Transcriptional plasticity in response to serum appeared to be species specific as the percentage of genes modulated was independent of the species abundance in the community.

**Fig 6.**
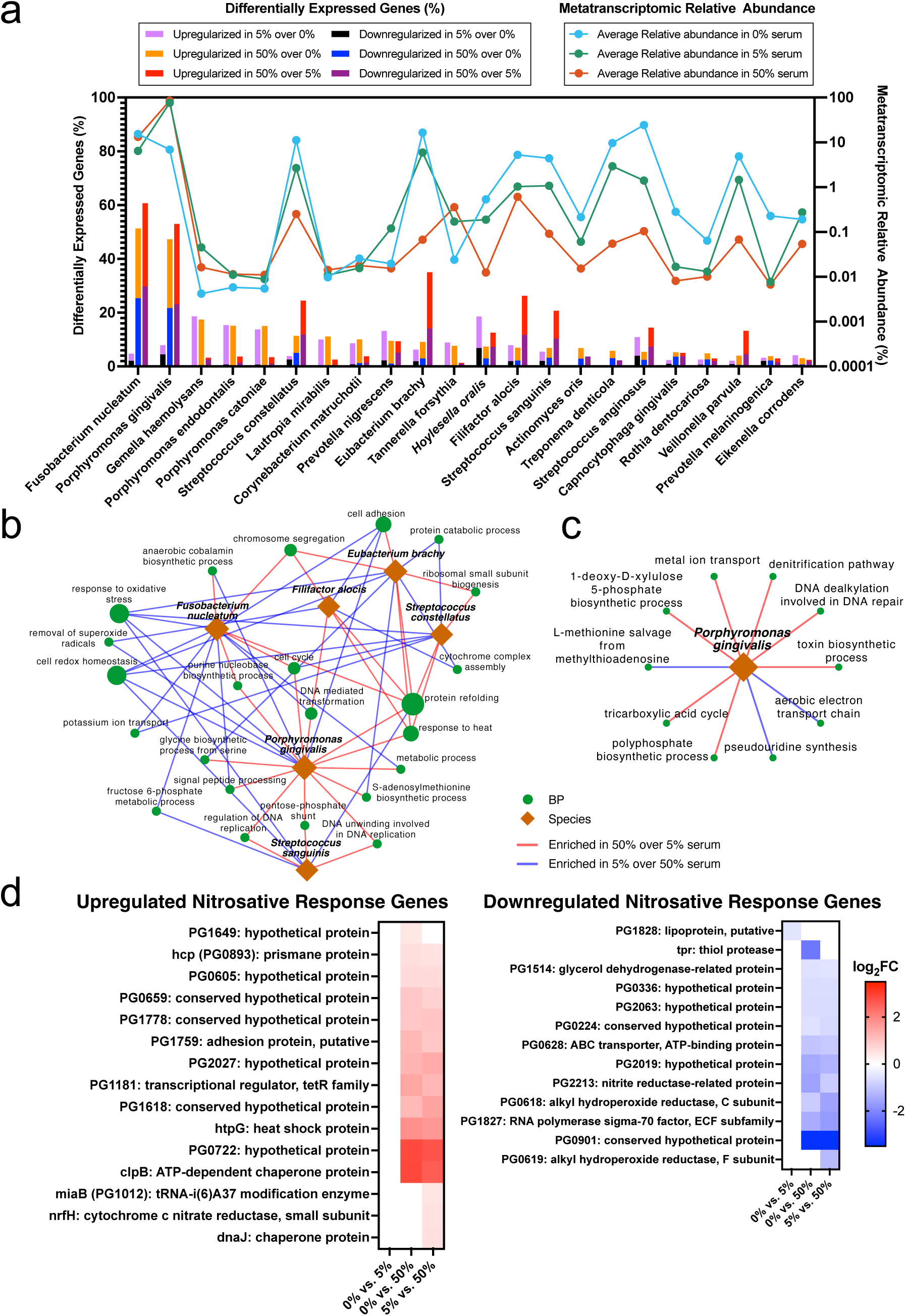
Changes in metatranscriptomic activities within species when growing as part of the 22 spp. community under different serum concentrations. (a) Bar plot showing the percentage of differentially expressed genes relative to the total number of genes detected per species, comparing steady states across serum concentrations. Lines represent the average metatranscriptomic relative abundance of each species across serum conditions. (b) Network of biological processes (BPs) enriched or depleted in 5% vs. 50% serum in the six species most responsive to serum. Diamonds represent species, and circles represent BPs, with edges connecting species and BPs enriched in either 5% (blue) or 50% serum (red). Only BPs with node degree > 2 are shown, with node size proportional to degree. (c) Network of BPs uniquely enriched or depleted in 5% vs. 50% serum in *Porphyromonas gingivalis*. (d) Heatmaps showing up- and down- regulated genes part of the response to nitrosative stress in *P. gingivalis*. The set of nitrosative stress response genes were identified by previous *in vitro* studies (Belvin et al 2019 [46], Lewis et al 2012 [47]). Differential expression across panels (a–d) was determined using DESeq2 (log2FC > 0.3785 or < –0.3785, FDR < 0.05).

We next analyzed the biological processes affected in the most serum-responsive species (Supplementary Fig. S7). The network in Fig. 6b illustrates the BPs enriched or depleted when comparing 5% and 50% serum across multiple species. Protein refolding was upregulated in 5 of the 6 species analyzed, confirming our findings of a community-wide response to maintain protein homeostasis at higher serum concentrations. Conversely, oxidative stress responses and cell redox homeostasis processes were downregulated, also confirming community-wide findings. Among other BPs downregulated in response to serum in several species was cell adhesion, with *P. gingivalis* downregulating its adhesion protein and virulence factor fimbrillin (*fimA*), which has been previously shown to decrease in response to serum and saliva [45]. One key BP downregulated in several species by higher serum was potassium transport. This function has been shown to be downregulated in human subgingival communities during periodontitis [18].

Distinct responses to higher serum occurred in individual species (Fig. S7, Supplementary Tables S6-11). For instance, *P. gingivalis* uniquely upregulated the denitrification pathway, which could serve as precursor for RNSs (Fig. 6c). *nrfH*, a membrane-bound cytochrome c which acts as a redox partner to the nitrite reductase *nrfA*, was upregulated. Additionally, species-specific analysis of *P. gingivalis* revealed upregulation of DNA repair (Fig. 6c). Given that *P. gingivalis* was the most abundant species in both 16S rRNA and metatranscriptomic analyses, this species may play a pivotal role in driving nitric oxide and RNS release.

Previous *in vitro* studies identified several genes in *P. gingivalis* as differentially regulated when growing in nitrite [46, 47]. Upon comparison, we found that a significant portion of nitrite-responsive genes previously identified were also differentially expressed in *P. gingivalis* in our community model in response to serum (Fig. 6d and Supplementary Fig. S8). This included upregulation of major nitrosative stress response genes *hcp* (PG0893), *nrfH*, and *clpB*. This strongly supports the hypothesis that nitrosative stress plays a key role in modulating transcriptomic activity in this community model under higher serum conditions.

In addition, the GO biological process S-adenosylmethionine biosynthesis was upregulated in *P. gingivalis*, with the gene *metK,* which encodes the enzyme responsible for S-adenosylmethionine (SAM) increased under higher serum in *P. gingivalis* and *F. alocis* (Supplementary Table S6 and S10). SAM has been shown to participate in a variety of cellular processes including DNA repair, but also cell growth, survival and quorum sensing [48–51]. The core species *F. nucleatum* showed a strong upregulation of nickel transport under 50% serum suggesting this metal ion is important for its survival in this environment. Meanwhile, the commensal *S. sanguinis* upregulated, under 50% serum, ABC transporters for glycine betaine, which acts as a protectant against osmotic stress in bacteria [52], suggesting the high protein levels under high serum may have created colloid osmotic pressures harmful for this species.

### Species-specific transcriptomic alterations in *P. gingivalis* explain increased pH under serum

Given the observed community-wide changes in KOs associated with increased ammonia production and reduced acidic byproducts, we investigated species-specific metabolic alterations in *P. gingivalis* and *F. nucleatum*, the two most abundant species in the community, also exhibiting significant changes in metatranscriptomic activity in response to serum. We focused on identifying significantly regulated genes that could potentially affect the environmental pH. Figure 7a shows that genes related to ammonia production, such as *pdxH*, *nrfH*, PG_0893 (*hcp*), and *ribD*, were upregulated in *P. gingivalis* in response to serum. In contrast, *F. nucleatum* exhibited upregulation of FN0488 (glutamate dehydrogenase), while genes involved in acid production, such as FN1162 (hydroxyacylglutathione hydrolase) and FN0814 (propionate CoA-transferase), were downregulated. These findings point to the specific reactions in these microorganisms capable of affecting the environmental pH, with *P. gingivalis* showing a plethora of ammonia producing reactions upregulated by serum.

**Fig 7.**
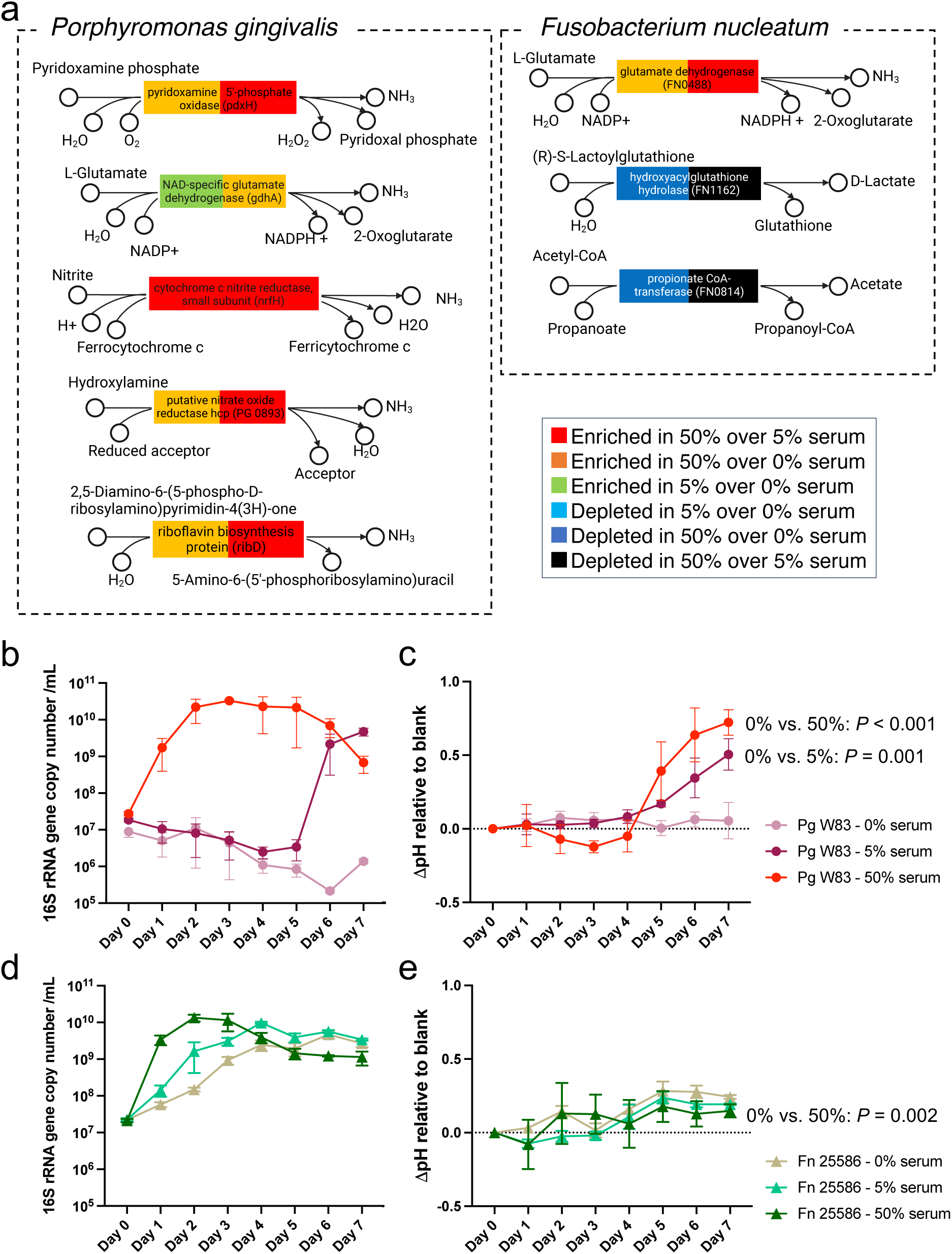
Effect of serum on the metabolic activities, growth and ability to modify the pH of *Porphyromonas gingivalis* and *Fusobacterium nucleatum*. (a) Differentially expressed genes involved in ammonia and acidic end-product formation in *P. gingivalis* and *F. nucleatum* as measured in steady-state chemostat cultures. (b) Biomass of *P. gingivalis* in batch monoculture under varying serum concentrations. (c) pH change (ΔpH) relative to fresh medium for *P. gingivalis* cultured in varying serum concentrations. (d) Biomass of *F. nucleatum* cultured under varying serum concentrations. (e) pH changes (ΔpH) relative to fresh medium for *F. nucleatum* cultures under varying serum concentrations. Statistical significance for ΔpH (panels c and e) was assessed on day 7 using one-way ANOVA with Bonferroni post hoc correction; only pairs showing significant differences were annotated.

To validate these transcriptomic findings, we conducted culture experiments to assess the growth of *P. gingivalis* and *F. nucleatum* under varying serum concentrations and to explore their influence on environmental pH. Fresh media incubated under the same conditions was included as the benchmark, and the difference between the culture pH and fresh media was denoted as ΔpH, which was used to evaluate the impact of the bacteria on environmental pH. As shown in Fig. 7b, *P. gingivalis* needed serum in the medium to grow as a monoculture, in contrast to its ability to grow as an abundant community member in the 22-species community with no serum (Fig. 2c). These results, unexpectedly, showed the essential role of inter-species cooperation to support growth of this pathogen in mucin. Fig. 7b also shows that 50% serum markedly decreased the lag phase of *P. gingivalis*, with these results agreeing with the advantage *P. gingivalis* gained in our chemostat community under serum. *F. nucleatum* also benefited from the presence of higher serum, although it was able to grow as a monoculture without it (Fig. 7d). With respect to the ability of these species to modify the pH, only *P. gingivalis*, when growing under serum, markedly increased the ΔpH, while *F. nucleatum* only slightly modified the pH towards alkalinity (Fig. 7c, e). Collectively, these findings suggest that *P. gingivalis* is a key driver of the metabolic adaptations observed in the microbial community, contributing significantly to the regulation of environmental pH under serum-enriched conditions.

## Discussion

In vitro model systems are essential for dissecting the ecological factors that shape microbiome communities [53–55]. Synthetic communities grown in chemostat-based continuous culture systems provide reproducible and tractable laboratory models that allow systematic evaluation of the influence of different ecological variables on community phenotypes. Here, we present a synthetic community comprised of prevalent and abundant human subgingival species which can be used as a model to probe how specific taxa, gene functions, and environmental factors shape community traits. The complementary use of omics analytical tools allows in-depth phenotypic characterization of communities over the longitudinal course of experiments. This model follows pioneering work in the pre-omics era that developed defined inocula oral communities in chemostats greatly advancing understanding of oral microbial ecology [15, 56]. The current model, however, includes a greater number of species, selected based on current knowledge of the subgingival microbiome, and utilizes omics techniques for in-depth community characterization [15, 56].

The community model was established using a mucin-based basal medium to mimic salivary fluid, the primary nutrient source for oral microorganisms. Prior work using this medium has shown that oral bacteria cooperate through complementary glycosidase and protease activities to degrade the mucin substrate, which is able to support growth of a polymicrobial community [57]. Subgingival communities, however, are also continuously bathed by GCF, which flows more rapidly and changes in composition under inflammatory conditions [22, 58]. We used serum as a surrogate as it resembles GCF [59]. Serum has been previously utilized as a sole nutrient source for native subgingival communities, with prior studies showing oral microbial consortia readily degrade serum glycoproteins [60]. Although both mucin and serum are rich in glycoproteins, these two nutritional substrates differ, as demonstrated here by the inability of *P. gingivalis* to grow in mucin as a monoculture, where it depended on the presence of the community, while addition of serum to the mucin medium readily supported monoculture growth of *P. gingivalis*. The preferential utilization of serum by *P. gingivalis* likely underlies its competitive dominance observed when serum became available.

The community model included species that are abundant members of the human subgingival microbiome in health as well as in periodontitis. Although some species became dominant, all species were transcriptionally active and therefore the model provides a platform to interrogate drivers of shifts in community configuration between healthy- and disease-like states. Our study used this community model to evaluate the influence of nutrients, specifically serum, on the community properties. We were, however, unable to evaluate the influence of serum on a health-like community as the community established in mucin, without serum, resembled a periodontitis-associated community with its composition dominated by a diverse group of core species and pathobionts, while health-associated species were present in low abundance. Although we supplied oxygen to the culture, the community readily metabolized it and maintained a very low redox potential, suppressing the growth of facultative health-associated commensals. These results suggest that the nutritional pressure of serum is not required to establish a periodontitis-like community, and other factors such as anaerobiosis may play a more important role to promote pathobiont growth and suppress health-associated species. Our results are comparable to those of Nagynte et al. [11] who enriched natural oral communities in a mucin-based medium, under anaerobiosis, also testing the effect of adding heat-inactivated serum on community composition, finding that the mucin-rich medium supported a community enriched for certain periodontitis-associated bacteria, such as *T. denticola*, with very low levels of health-associated commensals, while addition of serum favored the growth of other periodontitis-associated taxa, including *P. gingivalis* and *F. alocis* [11]. Similar to these results, health-associated commensals remained at a very low level in our communities, *T. denticola* was enriched in mucin without serum, and *P. gingivalis* and *F. alocis* were favored by serum.

Although serum appears as non-essential for a periodontitis-like community to establish, it changed the community in ways that recapitulate periodontitis-associated dysbiotic shifts, including promoting an increase in biomass [5, 6], increasing the environmental pH and changing gene expression in a manner that parallels some of the transcriptomic alterations seen in human periodontitis. Some of these transcriptomic changes included increased amino acid catabolism, especially upregulation of histidine degradation, as well as increased nitric oxidoreductase activity, chaperone and protein folding activities and downregulation of potassium uptake, which have been previously reported as associated with metatranscriptomes of periodontitis sites compared to health [18, 61]. Prior studies also found an increase in the GO term lipid A biosynthesis in periodontitis compared to health [18], with our model finding that serum upregulated lipopolysaccharide biosynthesis in a concentration-dependent manner (Supplementary Fig. S4). Lipopolysaccharides are critical virulence factors for induction of inflammation by the oral microbiome [62]. Furthermore, a study dissecting longitudinal microbiome transcriptomic changes during periodontitis progression recently found that activation of cobalamin contributes to progression [17], while we found serum increased biosynthesis of corrin, which forms the structural core of cobalamin. Altogether these results show that although serum as a nutrient source is not necessary for a community with periodontitis-like taxonomic configuration to establish, serum plays a role shaping the community properties increasing its dysbiotic-like features.

Serum was seen to promote a more basic environmental pH due to increased ammonia production and decreased acidic end-products. *P. gingivalis*, the most abundant species under serum, was seen to play a major role increasing the pH as seen from its transcriptomic profile and its ability to increase the pH in monoculture. Previous work has shown that *P. gingivalis* can only grow in conditions with pH values between 6.7 to 8.0 [63]. In addition, environmental pH is not only critical for the growth of *P. gingivalis* but also for its virulence as the enzymatic activity of gingipains, the potent cysteine proteinases of *P. gingivalis*, is optimal between a pH of 7.5 to 8.0 [64]. As seen here, *P. gingivalis*, like other microbes, modifies the environmental pH to a favorable one for its own growth. In our chemostat-grown community, we controlled the pH around neutrality, but our results suggest that in an uncontrolled environment the pH would have increased, potentially influencing the fitness and gene expression of *P. gingivalis* itself and other community members. Indeed, another abundant community member, *F. nucleatum*, can tolerate a wide range of pH values but when cultured at a pH above 8 its phenotype changes with cells becoming significantly elongated, increasing their surface hydrophobicity and forming biofilms [65]. Interestingly, we observed under high serum the presence of very elongated rods, possibly *F. nucleatum*, associated with planktonic microbial aggregates of increased size. Although, the pH of the whole vessel was controlled around neutrality, it is possible that pH gradients existed within these microbial aggregates in which cells exposed to higher pH values changed their phenotype becoming elongated and making them more prone to co-adhesion. The pH of the gingival crevice has been shown to increase from below neutrality in health to values greater than 8 in periodontitis [66], and therefore our results suggest that an increased flow of the serum-like inflammatory exudate in periodontitis and the community activities of *P. gingivalis* contribute to this environmental shift which may shape the community phenotype.

High serum induced protein and DNA repair mechanisms at a community-wide level and when evaluating transcriptional changes within species. By comparing the transcriptomic response of *P. gingivalis* to high serum in our community with previous in vitro investigations of the transcriptional response of monocultures of *P. gingivalis* to nitrite [46, 47], it was clear that there was a significant overlap, and therefore the stress response seen in the community likely resulted from nitrosative stress. Prior studies in humans show upregulation of nitric oxidoreductase activity and proteostasis response genes in periodontitis compared to health and it is therefore likely that in vivo, the serum-like inflammatory exudate is responsible for exerting this pressure on the microbiome [18]. RNS is also a likely stress encountered by oral microorganisms as they leave the mouth and translocate systemically through the circulation [67].

In conclusion, we present here a synthetic community maintained under a controlled environment in continuous culture in a mucin-rich medium serving as a model of the human subgingival microbiome, which can be used to interrogate the ecological variables that shape community configuration and function. Using this community, we explored the role of serum as an ecological modifier finding that serum increased the community biomass, promoted polymicrobial aggregate formation, increased the environmental pH and induced transcriptional activities that paralleled those of in vivo microbiome communities in periodontitis. Although mucin was sufficient to sustain a periodontitis-like community, addition of serum favored the outgrowth of *P. gingivalis* and modulated the community properties. The serum-like exudate that bathes subgingival communities serves therefore as a nutritional pressure that likely perpetuates dysbiosis.

## Data Availability Statement

The microbiome sequence data was deposited in the Sequence Reads Archive (accession number PRJNA1336263). Data that support the findings of this study are available from the authors upon reasonable request.

## Acknowledgements

These studies were funded by National Institutes of Health grants R21DE023967 (P.I.D), R21DE034093 (P.I.D), K99DE034829 (L.L), and the FONDECYT grant 1251739 (A.H) from ANID, the Chilean government. D.S. was supported by NIH grant K12DE027827.

## Notes

### Competing Interest Statement

The authors have declared no competing interest.

## References

1. Lloyd-Price J, Arze C, Ananthakrishnan AN, Schirmer M, Avila-Pacheco J, Poon TW et al. Multi-omics of the gut microbial ecosystem in inflammatory bowel diseases. Nature. 2019;569:655–62

2. Lee JY, Tsolis RM, Baumler AJ. The microbiome and gut homeostasis. Science. 2022;377:eabp9960

3. Usyk M, Carlson L, Schlecht NF, Sollecito CC, Grassi E, Wiek F et al. Cervicovaginal microbiome and natural history of chlamydia trachomatis in adolescents and young women. Cell. 2025;188:1051–61 e12

4. Abusleme L, Hoare A, Hong BY, Diaz PI. Microbial signatures of health, gingivitis, and periodontitis. Periodontol 2000. 2021;86:57–78

5. Diaz PI, Hoare A, Hong BY. Subgingival microbiome shifts and community dynamics in periodontal diseases. J Calif Dent Assoc. 2016;44:421–35

6. Abusleme L, Dupuy AK, Dutzan N, Silva N, Burleson JA, Strausbaugh LD et al. The subgingival microbiome in health and periodontitis and its relationship with community biomass and inflammation. ISME J. 2013;7:1016–25

7. Griffen AL, Beall CJ, Campbell JH, Firestone ND, Kumar PS, Yang ZK et al. Distinct and complex bacterial profiles in human periodontitis and health revealed by 16s pyrosequencing. ISME J. 2012;6:1176–85

8. Lamont RJ, Hajishengallis G. Polymicrobial synergy and dysbiosis in inflammatory disease. Trends Mol Med. 2015;21:172–83

9. Hajishengallis G. The inflammophilic character of the periodontitis-associated microbiota. Mol Oral Microbiol. 2014;29:248–57

10. Li L, Hayashi-Okada Y, Falkner KL, Shimizu Y, Zambon JJ, Kirkwood KL et al. Effect of an intensive antiplaque regimen on microbiome outcomes after nonsurgical periodontal therapy. J Periodontol. 2025;96:241–54

11. Naginyte M, Do T, Meade J, Devine DA, Marsh PD. Enrichment of periodontal pathogens from the biofilms of healthy adults. Sci Rep. 2019;9:5491

12. Baraniya D, Naginyte M, Chen T, Albandar JM, Chialastri SM, Devine DA et al. Modeling normal and dysbiotic subgingival microbiomes: Effect of nutrients. J Dent Res. 2020:22034520902452

13. Ter Steeg PF, Van der Hoeven JS, De Jong MH, Van Munster PJ, Jansen MJ. Enrichment of subgingival microflora on human serum leading to accumulation of bacteroides species, peptostreptococci and fusobacteria. Antonie Van Leeuwenhoek. 1987;53:261–72

14. Celis AI, Relman DA, Huang KC. The impact of iron and heme availability on the healthy human gut microbiome in vivo and in vitro. Cell Chem Biol. 2023;30:110–26 e3

15. Bradshaw DJ, Marsh PD, Watson GK, Allison C. Role of fusobacterium nucleatum and coaggregation in anaerobe survival in planktonic and biofilm oral microbial communities during aeration. Infect Immun. 1998;66:4729–32

16. Hoare A, Wang H, Meethil A, Abusleme L, Hong BY, Moutsopoulos NM et al. A cross-species interaction with a symbiotic commensal enables cell-density-dependent growth and in vivo virulence of an oral pathogen. ISME J. 2021;15:1490–504

17. Duran-Pinedo A, Solbiati JO, Teles F, Yanping Z, Frias-Lopez J. Longitudinal host-microbiome dynamics of metatranscription identify hallmarks of progression in periodontitis. Microbiome. 2025;13:119

18. Duran-Pinedo AE, Chen T, Teles R, Starr JR, Wang X, Krishnan K et al. Community-wide transcriptome of the oral microbiome in subjects with and without periodontitis. ISME J. 2014

19. Lamster IB. Evaluation of components of gingival crevicular fluid as diagnostic tests. Ann Periodontol. 1997;2:123–37

20. Romano F, Iaderosa G, Corana M, Perotto S, Baima G, Di Scipio F et al. Comparing ionic profile of gingival crevicular fluid and saliva as distinctive signature of severe periodontitis. Biomedicines. 2022;10

21. Cuevas-Gonzalez MV, Cuevas-Gonzalez JC, Espinosa-Cristobal LF, Tovar-Carrillo KL, Saucedo-Acuna RA, Garcia-Calderon AG et al. The potential of gingival crevicular fluid as a tool for molecular diagnosis: A systematic review. Biomed Res Int. 2024;2024:5560866

22. Ozkavaf A, Aras H, Huri CB, Mottaghian-Dini F, Tozum TF, Etikan I et al. Relationship between the quantity of gingival crevicular fluid and clinical periodontal status. Journal of oral science. 2000;42:231–8

23. Hatipoglu H, Yamalik N, Berberoglu A, Eratalay K. Impact of the distinct sampling area on volumetric features of gingival crevicular fluid. Journal of periodontology. 2007;78:705–15

24. Kinniment SL, Wimpenny JW, Adams D, Marsh PD. Development of a steady-state oral microbial biofilm community using the constant-depth film ferrnenter. Microbiology. 1996;142:631–38

25. Schincaglia GP, Hong BY, Rosania A, Barasz J, Thompson A, Sobue T et al. Clinical, immune, and microbiome traits of gingivitis and peri-implant mucositis. J Dent Res. 2017;96:47–55

26. Schloss PD, Westcott SL, Ryabin T, Hall JR, Hartmann M, Hollister EB et al. Introducing mothur: Open-source, platform-independent, community-supported software for describing and comparing microbial communities. Appl Environ Microbiol. 2009;75:7537–41

27. Edgar RC, Haas BJ, Clemente JC, Quince C, Knight R. Uchime improves sensitivity and speed of chimera detection. Bioinformatics. 2011;27:2194–200

28. Dewhirst FE, Chen T, Izard J, Paster BJ, Tanner AC, Yu WH et al. The human oral microbiome. J Bacteriol. 2010;192:5002–17

29. Huttenhower Lab HU. https://huttenhower.sph.harvard.edu/kneaddata

30. Hattori M. Finishing the euchromatic sequence of the human genome. Tanpakushitsu Kakusan Koso. 2005;50:162–8

31. Langmead B, Salzberg SL. Fast gapped-read alignment with bowtie 2. Nat Methods. 2012;9:357–9

32. Li H, Handsaker B, Wysoker A, Fennell T, Ruan J, Homer N et al. The sequence alignment/map format and samtools. Bioinformatics. 2009;25:2078–9

33. Buchfink B, Reuter K, Drost HG. Sensitive protein alignments at tree-of-life scale using diamond. Nat Methods. 2021;18:366–68

34. Ogata H, Goto S, Sato K, Fujibuchi W, Bono H, Kanehisa M. Kegg: Kyoto encyclopedia of genes and genomes. Nucleic Acids Res. 1999;27:29–34

35. Ashburner M, Ball CA, Blake JA, Botstein D, Butler H, Cherry JM et al. Gene ontology: Tool for the unification of biology. The gene ontology consortium. Nat Genet. 2000;25:25–9

36. Love MI, Huber W, Anders S. Moderated estimation of fold change and dispersion for rna-seq data with deseq2. Genome Biol. 2014;15:550

37. Supek F, Bošnjak M, Škunca N, Šmuc T. Revigo summarizes and visualizes long lists of gene ontology terms. PLoS One. 2011;6:e21800

38. Katajamaa M, Miettinen J, Oresic M. Mzmine: Toolbox for processing and visualization of mass spectrometry based molecular profile data. Bioinformatics. 2006;22:634–6

39. Wishart DS, Guo A, Oler E, Wang F, Anjum A, Peters H et al. Hmdb 5.0: The human metabolome database for 2022. Nucleic Acids Res. 2022;50:D622–d31

40. Pang Z, Zhou G, Ewald J, Chang L, Hacariz O, Basu N et al. Using metaboanalyst 5.0 for lc-hrms spectra processing, multi-omics integration and covariate adjustment of global metabolomics data. Nat Protoc. 2022;17:1735–61

41. Diaz PI, Hoare A, Hong B-Y. Subgingival microbiome shifts and community dynamics in periodontal diseases. Journal of the California Dental Association. 2016;44:421–35

42. Simon-Soro A, Ren Z, Krom BP, Hoogenkamp MA, Cabello-Yeves PJ, Daniel SG et al. Polymicrobial aggregates in human saliva build the oral biofilm. MBio. 2022;13:e00131–22

43. Gladwin MT, Shelhamer JH, Schechter AN, Pease-Fye ME, Waclawiw MA, Panza JA et al. Role of circulating nitrite and s-nitrosohemoglobin in the regulation of regional blood flow in humans. Proc Natl Acad Sci U S A. 2000;97:11482–7

44. Giovannoni G, Land JM, Keir G, Thompson EJ, Heales SJ. Adaptation of the nitrate reductase and griess reaction methods for the measurement of serum nitrate plus nitrite levels. Ann Clin Biochem. 1997;34 (Pt 2):193–8

45. Xie H, Cai S, Lamont RJ. Environmental regulation of fimbrial gene expression in porphyromonas gingivalis. Infection and Immunity. 1997;65:2265–71

46. Belvin BR, Gui Q, Hutcherson JA, Lewis JP. The porphyromonas gingivalis hybrid cluster protein hcp is required for growth with nitrite and survival with host cells. Infection and Immunity. 2019;87:10.1128/iai.00572-18

47. Lewis JP, Yanamandra SS, Anaya-Bergman C. Hcpr of porphyromonas gingivalis is required for growth under nitrosative stress and survival within host cells. Infection and Immunity. 2012;80:3319–31

48. Ishiguro K, Arai T, Suzuki T. Depletion of s-adenosylmethionine impacts on ribosome biogenesis through hypomodification of a single rrna methylation. Nucleic Acids Research. 2019;47:4226–39

49. Wang SC, Frey PA. S-adenosylmethionine as an oxidant: The radical sam superfamily. Trends in Biochemical Sciences. 2007;32:101–10

50. Hanzelka BL, Greenberg E. Quorum sensing in vibrio fischeri: Evidence that s-adenosylmethionine is the amino acid substrate for autoinducer synthesis. J Bacteriol. 1996;178:5291–94

51. Schauder S, Shokat K, Surette MG, Bassler BL. The luxs family of bacterial autoinducers: Biosynthesis of a novel quorum-sensing signal molecule. Mol Microbiol. 2001;41:463–76

52. Van Der Heide T, Poolman B. Osmoregulated abc-transport system of lactococcus lactis senses water stress via changes in the physical state of the membrane. Proc Natl Acad Sci U S A. 2000;97:7102–06

53. Bradshaw DJ, Marsh PD, Hodgson RJ, Visser JM. Effects of glucose and fluoride on competition and metabolism within in vitro dental bacterial communities and biofilms. Caries Res. 2002;36:81–6

54. Weiss AS, Niedermeier LS, von Strempel A, Burrichter AG, Ring D, Meng C et al. Nutritional and host environments determine community ecology and keystone species in a synthetic gut bacterial community. Nat Commun. 2023;14:4780

55. Goldman DA, Xue KS, Parrott AB, Lopez JA, Vila JCC, Jeeda RR et al. Competition for shared resources increases dependence on initial population size during coalescence of gut microbial communities. Proc Natl Acad Sci U S A. 2025;122:e2322440122

56. Bradshaw DJ, Marsh PD, Allison C, Schilling KM. Effect of oxygen, inoculum composition and flow rate on development of mixed-culture oral biofilms. Microbiology. 1996;142 (Pt 3):623–9

57. Bradshaw DJ, Homer KA, Marsh PD, Beighton D. Metabolic cooperation in oral microbial communities during growth on mucin. Microbiology. 1994;140 (Pt 12):3407–12

58. Huynh AH, Veith PD, McGregor NR, Adams GG, Chen D, Reynolds EC et al. Gingival crevicular fluid proteomes in health, gingivitis and chronic periodontitis. Journal of periodontal research. 2014

59. Khurshid Z, Mali M, Naseem M, Najeeb S, Zafar MS. Human gingival crevicular fluids (gcf) proteomics: An overview. Dent J (Basel). 2017;5

60. Ter Steeg PF, Van der Hoeven JS, De Jong MH, Van Munster PJJ, Jansen MJH. Modelling the gingival pocket by enrichment of subgingival microflora in human serum in chemostats. Microb Ecol Health Dis. 1988;1:73–84

61. Jorth P, Turner KH, Gumus P, Nizam N, Buduneli N, Whiteley M. Metatranscriptomics of the human oral microbiome during health and disease. MBio. 2014;5:e01012–14

62. Dixon DR, Darveau RP. Lipopolysaccharide heterogeneity: Innate host responses to bacterial modification of lipid a structure. J Dent Res. 2005;84:584–95

63. McDermid AS, McKee AS, Marsh PD. Effect of environmental ph on enzyme activity and growth of bacteroides gingivalis w50. Infect Immun. 1988;56:1096–100

64. Takahashi N, Schachtele CF. Effect of ph on the growth and proteolytic activity of porphyromonas gingivalis and bacteroides intermedius. J Dent Res. 1990;69:1266–9

65. Zilm PS, Rogers AH. Co-adhesion and biofilm formation by fusobacterium nucleatum in response to growth ph. Anaerobe. 2007;13:146–52

66. Bickel M, Cimasoni G. The ph of human crevicular fluid measured by a new microanalytical technique. J Periodontal Res. 1985;20:35–40

67. Han YW, Wang X. Mobile microbiome: Oral bacteria in extra-oral infections and inflammation. J Dent Res. 2013;92:485–91

