## Supplementary Figures for "The Impact of Serum on a Complex Synthetic Community Model of the Subgingival Microbiome"

**Figure S1. Comparison of steady-state biomass in a five-species community cultured in continuous mode at two different dilution rates.** Growth conditions were 5% serum, continuous culture, anaerobic atmosphere (5% $CO_2$ in $N_2$), 37 °C, and pH 7.15 ± 0.15. Protocol 1 (open bars): D = 0.103 $h^{-1}$, Td = 6.72 h, F = 51.56 mL/h. Protocol 2 (dashed bars): D = 0.0462 $h^{-1}$, Td = 15 h, F = 23.21 mL/h. Biomass of *A. oris*, *S. sanguinis*, *F. nucleatum*, and *V. parvula* was assessed by colony-forming unit (CFU) counts per mL, whereas *P. gingivalis* abundance was determined determined as 16S rRNA gene copies per mL by qPCR. Differences in biomass between the two protocols were evaluated using t-test with the Bonferroni method for multiple-testing correction.

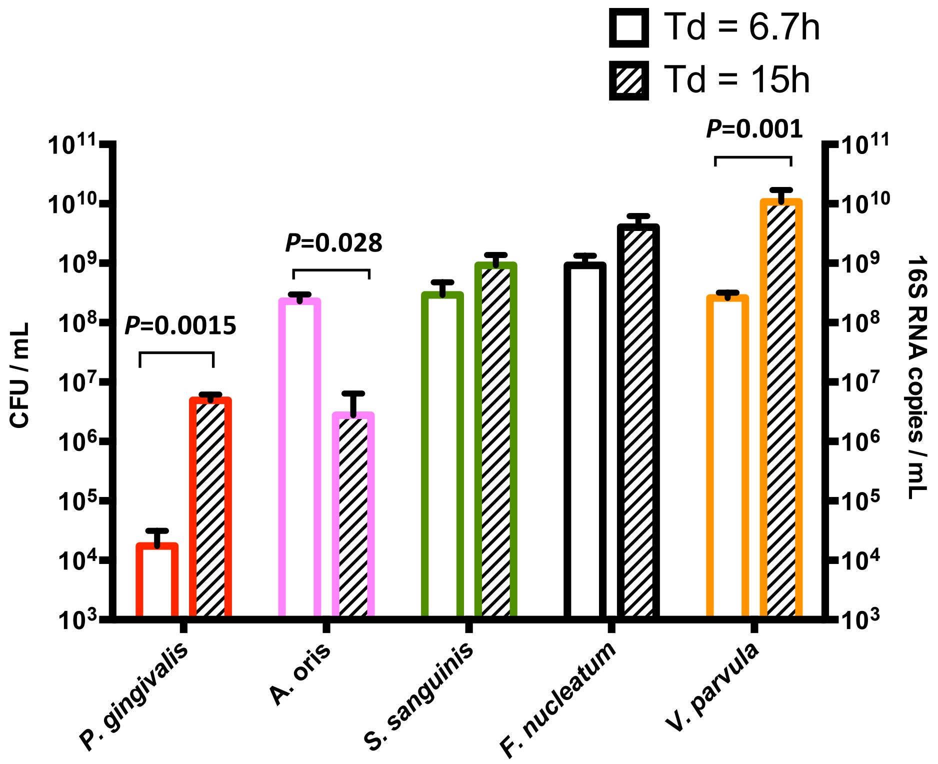

**Figure S2. Replicate analysis of community taxonomic composition under serum shift conditions.** Community composition during the assembly and theoretical steady-state phases (15–21 mean generation times, MGT) was characterized by 16S rRNA gene sequencing in two independent runs: run 1 (a) and run 2 (b). Both runs were initiated in 5% serum and subsequently shifted to 50% serum. (c) Spearman's correlations of the relative abundance of individual species between run 1 and run 2 across the full experimental period. (d) Principal coordinate analysis (PCoA) of Bray–Curtis beta-diversity of steady-state communities at different serum concentrations in run 1 and run 2. Significance of group differences was determined using PERMANOVA test with Bonferroni correction as post hoc analysis. There was no significant difference ($P > 0.05$) between the two runs within the same serum concentration.

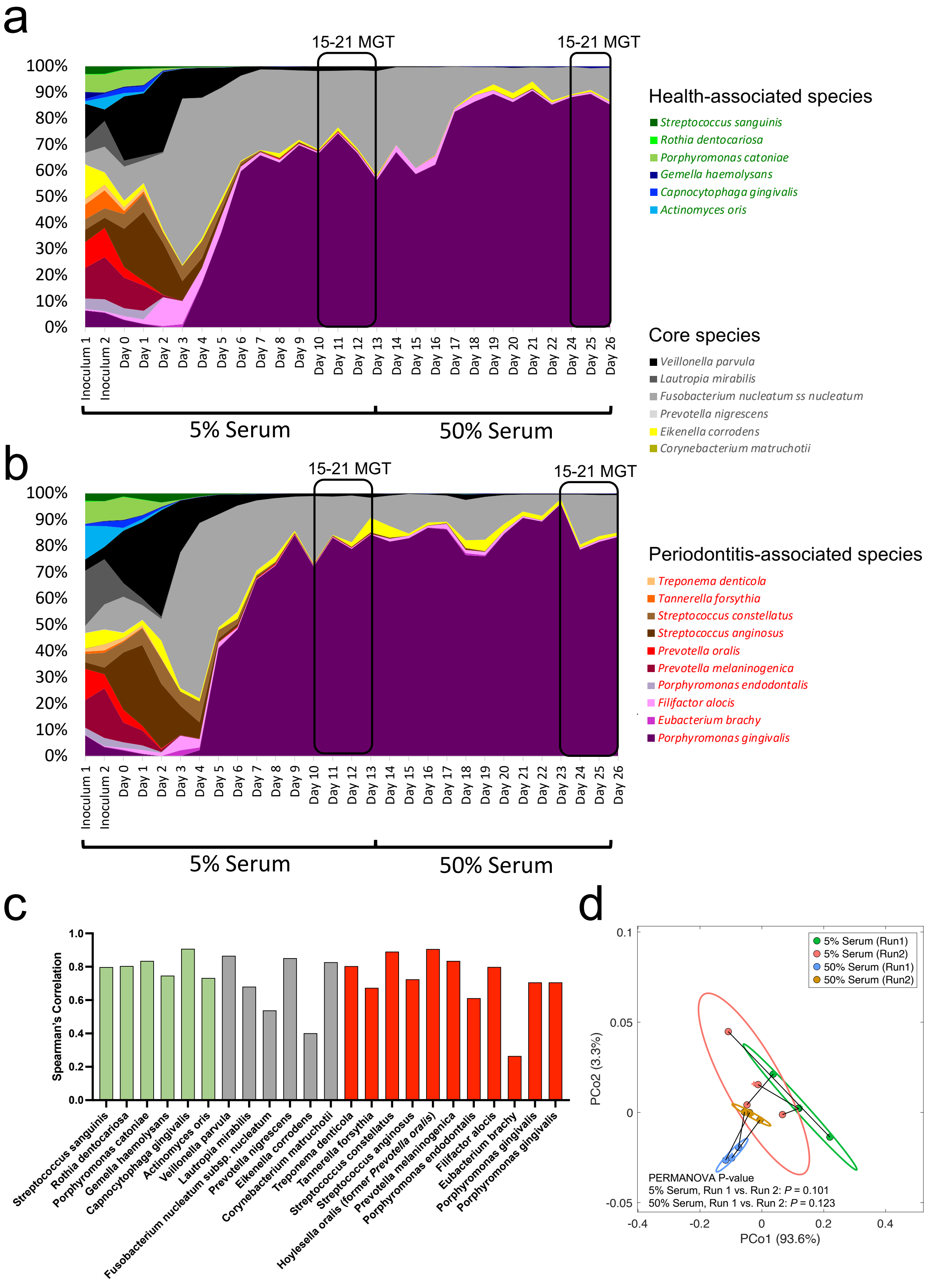
a
15-21 MGT
15-21 MGT
100%
90%
80%
70%
60%
50%
40%
30%
20%
10%
0%
Inoculum 1
Inoculum 2
Day 0
Day 1
Day 2
Day 3
Day 4
Day 5
Day 6
Day 7
Day 8
Day 9
Day 10
Day 11
Day 12
Day 13
Day 14
Day 15
Day 16
Day 17
Day 18
Day 19
Day 20
Day 21
Day 22
Day 24
Day 25
Day 26
5% Serum
50% Serum
Health-associated species
Streptococcus sanguinis
Rothia dentocariosa
Porphyromonas catoniae
Gemella haemolysans
Capnocytophaga gingivalis
Actinomyces oris
Core species
Veillonella parvula
Lautropia mirabilis
Fusobacterium nucleatum ss nucleatum
Prevotella nigrescens
Eikenella corrodens
Corynebacterium matruchotii
b
15-21 MGT
15-21 MGT
100%
90%
80%
70%
60%
50%
40%
30%
20%
10%
0%
Inoculum 1
Inoculum 2
Day 0
Day 1
Day 2
Day 3
Day 4
Day 5
Day 6
Day 7
Day 8
Day 9
Day 10
Day 11
Day 12
Day 13
Day 14
Day 15
Day 16
Day 17
Day 18
Day 19
Day 20
Day 21
Day 22
Day 23
Day 24
Day 25
Day 26
5% Serum
50% Serum
Periodontitis-associated species
Treponema denticola
Tannerella forsythia
Streptococcus constellatus
Streptococcus anginosus
Prevotella oralis
Prevotella melaninogenica
Porphyromonas endodontalis
Filifactor alocis
Eubacterium brachy
Porphyromonas gingivalis
c
Spearman's Correlation
1.0
0.8
0.6
0.4
0.2
0.0
Streptococcus sanguinis
Rothia dentocariosa
Porphyromonas catoniae
Gemella haemolysans
Capnocytophaga gingivalis
Actinomyces oris
Veillonella parvula
Lautropia mirabilis
Fusobacterium nucleatum subsp. nucleatum
Prevotella nigrescens
Eikenella corrodens
Corynebacterium matruchotii
Treponema denticola
Tannerella forsythia
Streptococcus constellatus
Streptococcus anginosus
Hoylesella oralis (former Prevotella oralis)
Prevotella melaninogenica
Porphyromonas endodontalis
Filifactor alocis
Eubacterium brachy
Porphyromonas gingivalis
Porphyromonas gingivalis
d
5% Serum (Run1)
5% Serum (Run2)
50% Serum (Run1)
50% Serum (Run2)
0.1
0.05
0
-0.05
PCo2 (3.3%)
-0.4
-0.2
0
0.2
0.4
PCo1 (93.6%)
PERMANOVA P-value
5% Serum, Run 1 vs. Run 2: P = 0.101
50% Serum, Run 1 vs. Run 2: P = 0.123

**Figure S3. Effect of serum on community evenness during the assembly period and theoretical steady state phases** (15-21 mean generation times, MGT) as characterized via 16S rRNA sequencing. (a–c) Daily evenness measured by the Shannon index over time under different serum conditions. (d) Bar plot comparing Shannon evenness at steady states across serum concentrations. Statistical significance was assessed using one-way ANOVA with Bonferroni correction for multiple comparisons.

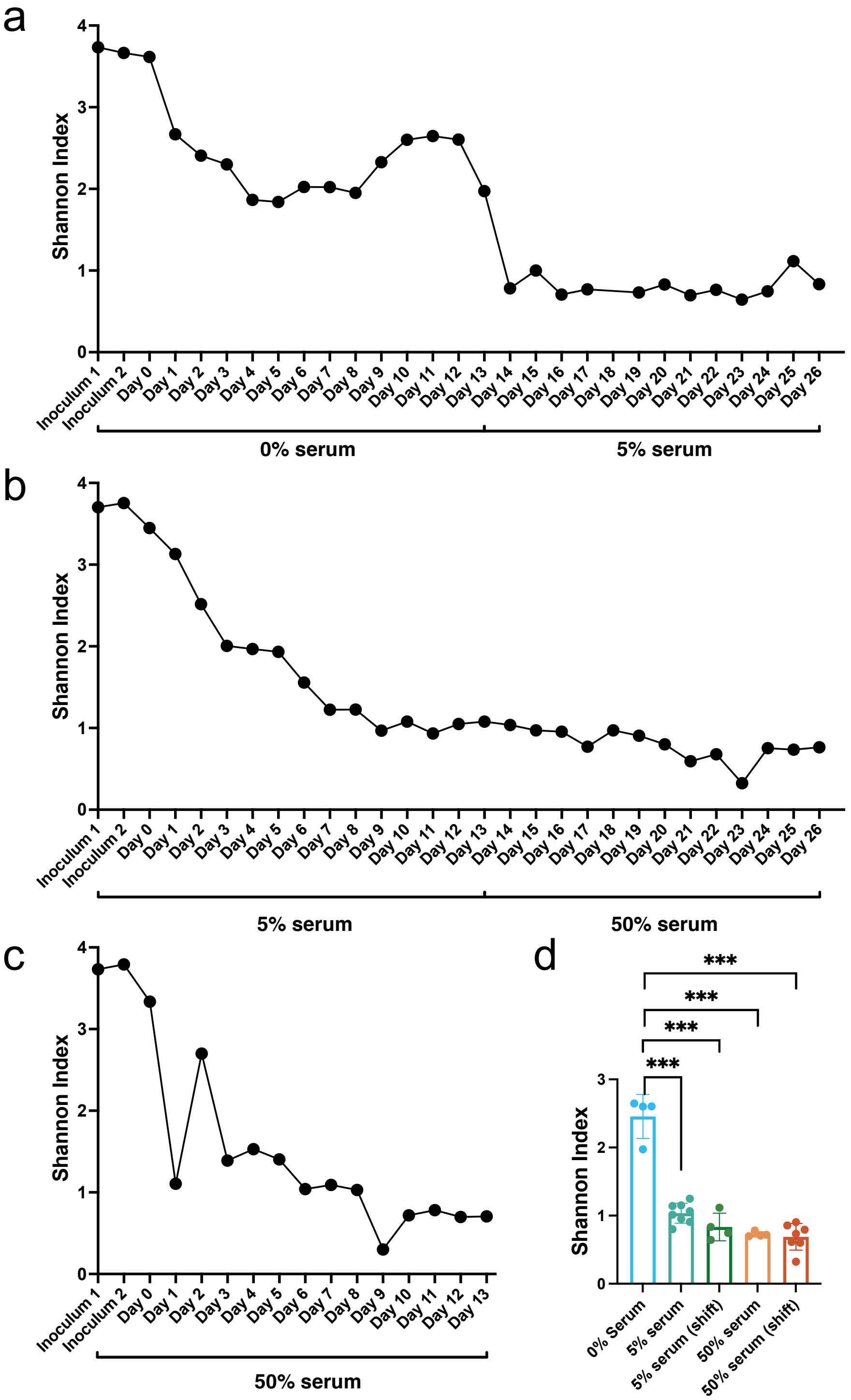

**Figure S4. Heatmap plot depicts log2 fold-changes (log2FC) of differentially expressed KEGG pathways across serum concentrations, identified by DESeq2.** Differential expression was filtered using log2FC > 0.3785 or < –0.3785 and FDR < 0.05.

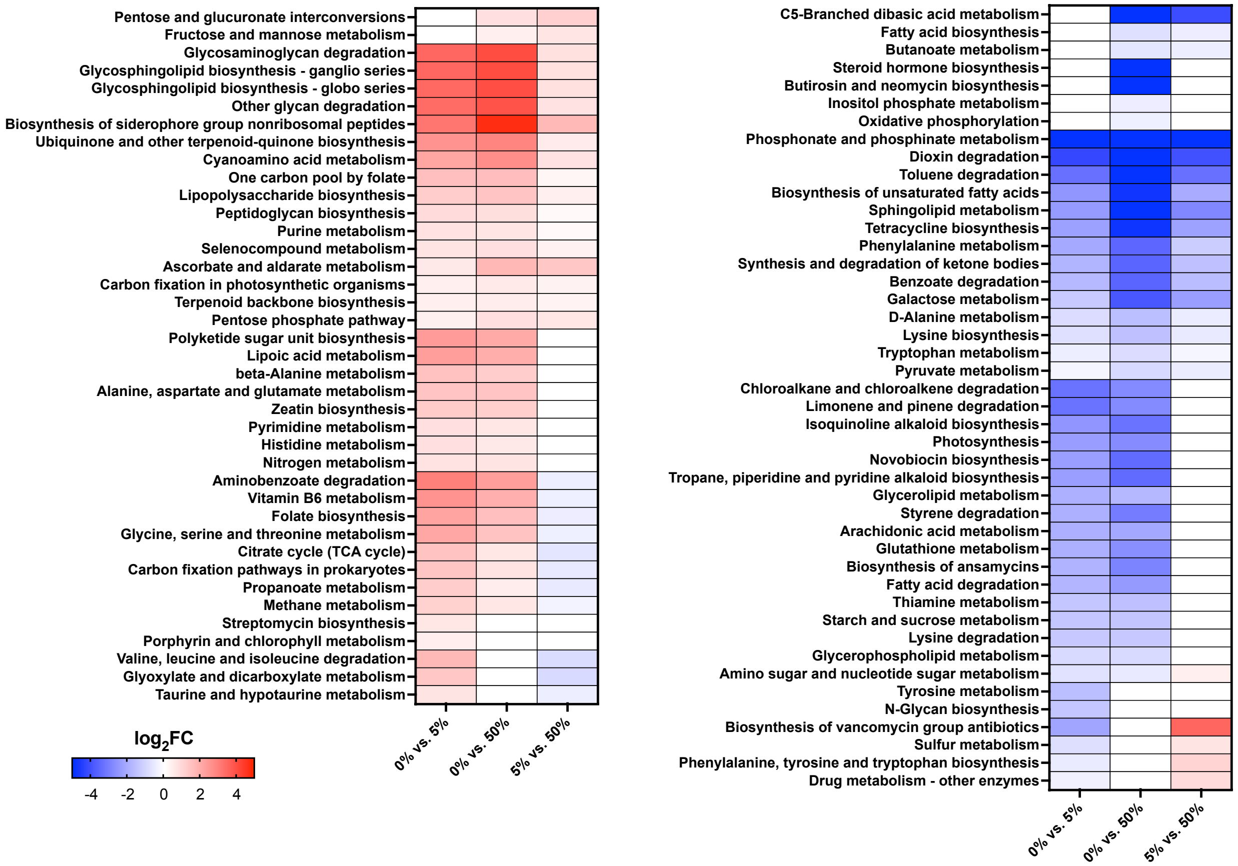

**Figure S5**. **Differential expression of UniRef90 gene families involved in oxidative stress across serum concentrations.** Heatmap showing log2 fold-changes (log2FC) of differentially expressed UniRef90 gene families related to the response to oxidative stress when comparing communities grown in 0%, 5%, and 50% serum. Differential expression was determined using DESeq2, with thresholds of log2FC > 0.3785 or < –0.3785 and FDR < 0.05.

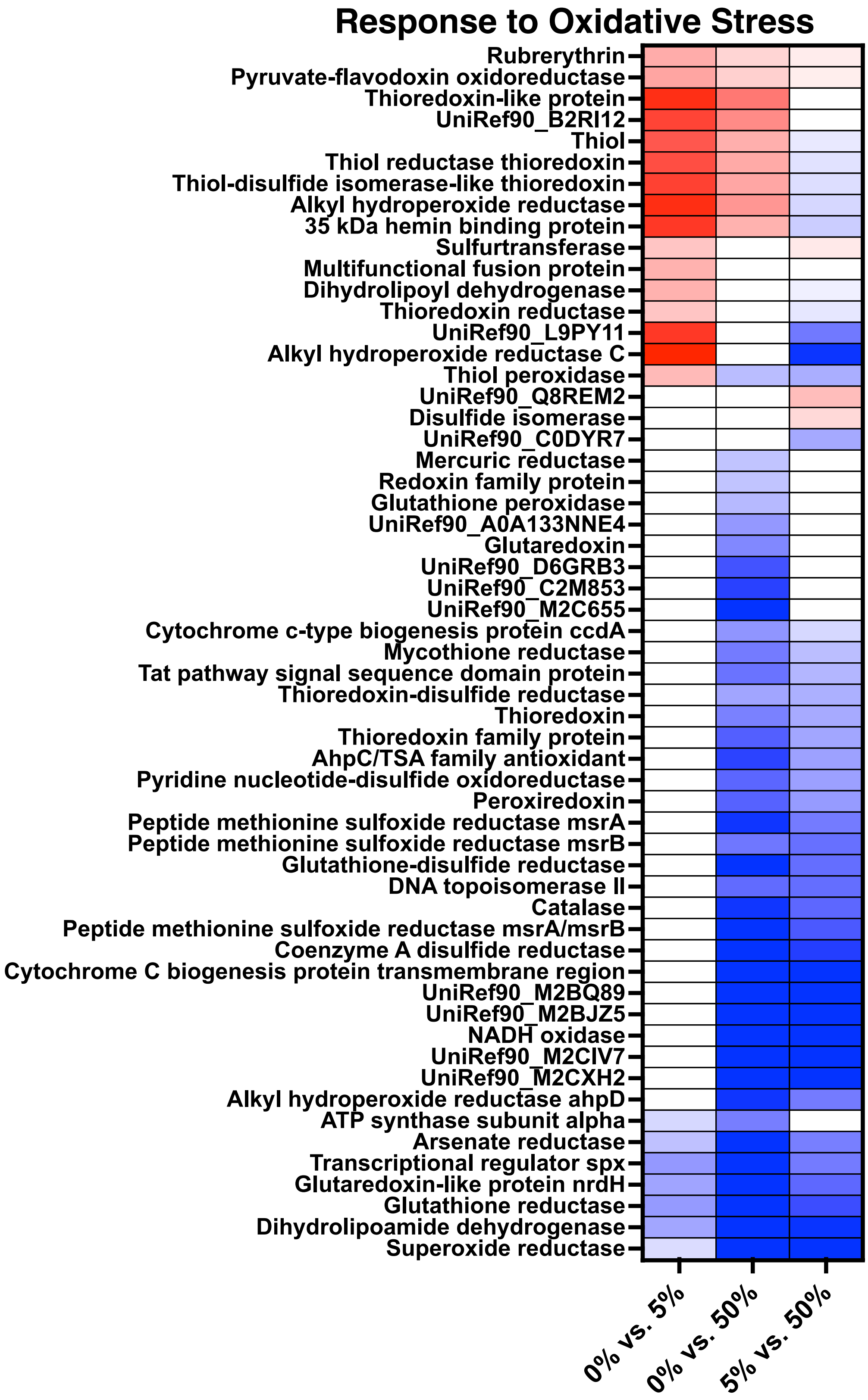
Response to Oxidative Stress
Rubrerythrin
Pyruvate-flavodoxin oxidoreductase
Thioredoxin-like protein
UniRef90_B2RI12
Thiol
Thiol reductase thioredoxin
Thiol-disulfide isomerase-like thioredoxin
Alkyl hydroperoxide reductase
35 kDa hemin binding protein
Sulfurtransferase
Multifunctional fusion protein
Dihydrolipoyl dehydrogenase
Thioredoxin reductase
UniRef90_L9PY11
Alkyl hydroperoxide reductase C
Thiol peroxidase
UniRef90_Q8REM2
Disulfide isomerase
UniRef90_C0DYR7
Mercuric reductase
Redoxin family protein
Glutathione peroxidase
UniRef90_A0A133NNE4
Glutaredoxin
UniRef90_D6GRB3
UniRef90_C2M853
UniRef90_M2C655
Cytochrome c-type biogenesis protein ccdA
Mycothione reductase
Tat pathway signal sequence domain protein
Thioredoxin-disulfide reductase
Thioredoxin
Thioredoxin family protein
AhpC/TSA family antioxidant
Pyridine nucleotide-disulfide oxidoreductase
Peroxiredoxin
Peptide methionine sulfoxide reductase msrA
Peptide methionine sulfoxide reductase msrB
Glutathione-disulfide reductase
DNA topoisomerase II
Catalase
Peptide methionine sulfoxide reductase msrA/msrB
Coenzyme A disulfide reductase
Cytochrome C biogenesis protein transmembrane region
UniRef90_M2BQ89
UniRef90_M2BJZ5
NADH oxidase
UniRef90_M2CIV7
UniRef90_M2CXH2
Alkyl hydroperoxide reductase ahpD
ATP synthase subunit alpha
Arsenate reductase
Transcriptional regulator spx
Glutaredoxin-like protein nrdH
Glutathione reductase
Dihydrolipoamide dehydrogenase
Superoxide reductase
0% vs. 5%
0% vs. 50%
5% vs. 50%

**Figure S6. Changes in the oxidation-reduction potential of steady states in different serum concentrations.** ANOVA tests were used to evaluate group-level differences with the Bonferroni method for multiple-comparison correction (***$P<0.001$).

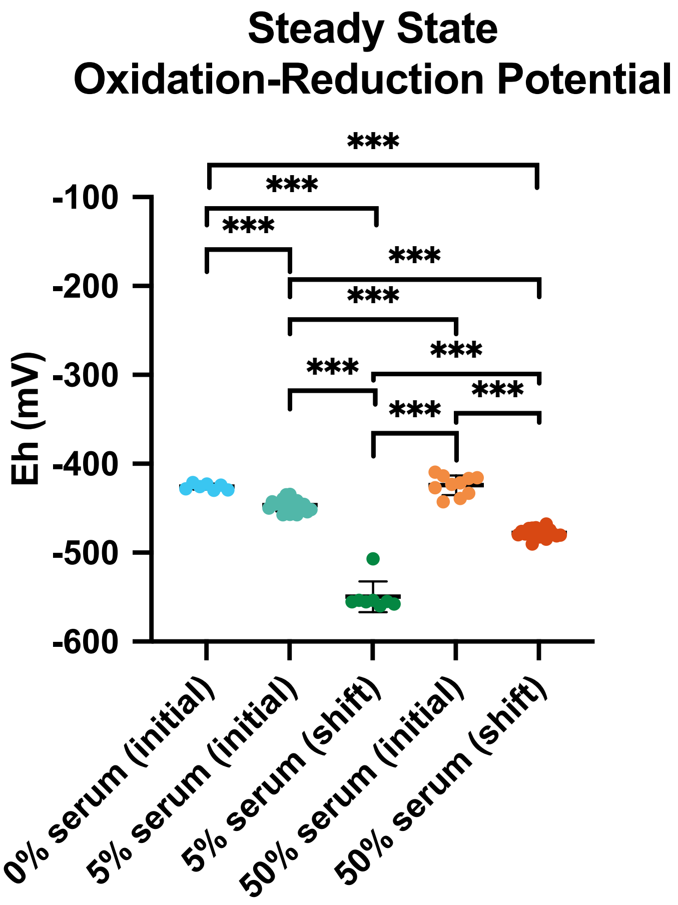

**Figure S7. Heatmaps summarize species-specific transcriptomic responses to serum.** The plots depict log2 fold-changes (log2FC) of differentially expressed biological processes (BPs) across serum concentrations, identified by DESeq2 from six species with the greatest response to serum, each with more than 20% of genes differentially expressed in any comparison. Differential expression was filtered using log2FC > 0.3785 or < –0.3785 and FDR < 0.05, with BPs unique to each species highlighted.

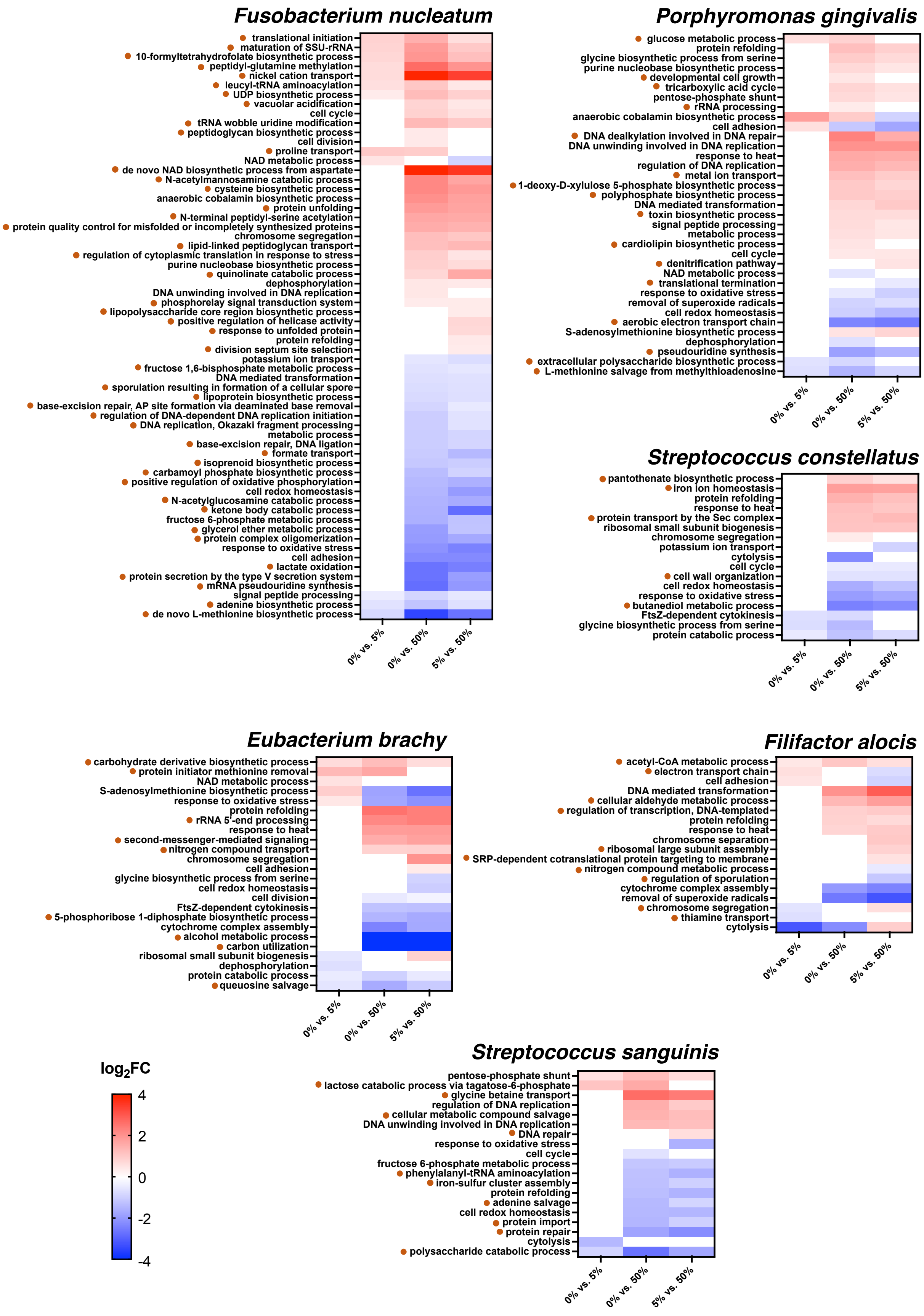
Fusobacterium nucleatum
translational initiation
maturation of SSU-rRNA
10-formyltetrahydrofolate biosynthetic process
peptidyl-glutamine methylation
nickel cation transport
leucyl-tRNA aminoacylation
UDP biosynthetic process
vacuolar acidification
cell cycle
tRNA wobble uridine modification
peptidoglycan biosynthetic process
cell division
proline transport
NAD metabolic process
de novo NAD biosynthetic process from aspartate
N-acetylmannosamine catabolic process
cysteine biosynthetic process
anaerobic cobalamin biosynthetic process
protein unfolding
N-terminal peptidyl-serine acetylation
protein quality control for misfolded or incompletely synthesized proteins
chromosome segregation
lipid-linked peptidoglycan transport
regulation of cytoplasmic translation in response to stress
purine nucleobase biosynthetic process
quinolinate catabolic process
dephosphorylation
DNA unwinding involved in DNA replication
phosphorelay signal transduction system
lipopolysaccharide core region biosynthetic process
positive regulation of helicase activity
response to unfolded protein
protein refolding
division septum site selection
potassium ion transport
fructose 1,6-bisphosphate metabolic process
DNA mediated transformation
sporulation resulting in formation of a cellular spore
lipoprotein biosynthetic process
base-excision repair, AP site formation via deaminated base removal
regulation of DNA-dependent DNA replication initiation
DNA replication, Okazaki fragment processing
metabolic process
base-excision repair, DNA ligation
formate transport
isoprenoid biosynthetic process
carbamoyl phosphate biosynthetic process
positive regulation of oxidative phosphorylation
cell redox homeostasis
N-acetylglucosamine catabolic process
ketone body catabolic process
fructose 6-phosphate metabolic process
glycerol ether metabolic process
protein complex oligomerization
response to oxidative stress
cell adhesion
lactate oxidation
protein secretion by the type V secretion system
mRNA pseudouridine synthesis
signal peptide processing
adenine biosynthetic process
de novo L-methionine biosynthetic process
0% vs. 5%
0% vs. 50%
5% vs. 50%
Porphyromonas gingivalis
glucose metabolic process
protein refolding
glycine biosynthetic process from serine
purine nucleobase biosynthetic process
developmental cell growth
tricarboxylic acid cycle
pentose-phosphate shunt
rRNA processing
anaerobic cobalamin biosynthetic process
cell adhesion
DNA dealkylation involved in DNA repair
DNA unwinding involved in DNA replication
response to heat
regulation of DNA replication
metal ion transport
1-deoxy-D-xylulose 5-phosphate biosynthetic process
polyphosphate biosynthetic process
DNA mediated transformation
toxin biosynthetic process
signal peptide processing
metabolic process
cardiolipin biosynthetic process
cell cycle
denitrification pathway
NAD metabolic process
translational termination
response to oxidative stress
removal of superoxide radicals
cell redox homeostasis
aerobic electron transport chain
S-adenosylmethionine biosynthetic process
dephosphorylation
pseudouridine synthesis
extracellular polysaccharide biosynthetic process
L-methionine salvage from methylthioadenosine
0% vs. 5%
0% vs. 50%
5% vs. 50%
Streptococcus constellatus
pantothenate biosynthetic process
iron ion homeostasis
protein refolding
response to heat
protein transport by the Sec complex
ribosomal small subunit biogenesis
chromosome segregation
potassium ion transport
cytolysis
cell cycle
cell wall organization
cell redox homeostasis
response to oxidative stress
butanediol metabolic process
FtsZ-dependent cytokinesis
glycine biosynthetic process from serine
protein catabolic process
0% vs. 5%
0% vs. 50%
5% vs. 50%
Eubacterium brachy
carbohydrate derivative biosynthetic process
protein initiator methionine removal
NAD metabolic process
S-adenosylmethionine biosynthetic process
response to oxidative stress
protein refolding
rRNA 5'-end processing
response to heat
second-messenger-mediated signaling
nitrogen compound transport
chromosome segregation
cell adhesion
glycine biosynthetic process from serine
cell redox homeostasis
cell division
FtsZ-dependent cytokinesis
5-phosphoribose 1-diphosphate biosynthetic process
cytochrome complex assembly
alcohol metabolic process
carbon utilization
ribosomal small subunit biogenesis
dephosphorylation
protein catabolic process
queuosine salvage
0% vs. 5%
0% vs. 50%
5% vs. 50%
Filifactor alocis
acetyl-CoA metabolic process
electron transport chain
cell adhesion
DNA mediated transformation
cellular aldehyde metabolic process
regulation of transcription, DNA-templated
protein refolding
response to heat
chromosome separation
ribosomal large subunit assembly
SRP-dependent cotranslational protein targeting to membrane
nitrogen compound metabolic process
regulation of sporulation
cytochrome complex assembly
removal of superoxide radicals
chromosome segregation
thiamine transport
cytolysis
0% vs. 5%
0% vs. 50%
5% vs. 50%
log2FC
4
2
0
-2
-4
Streptococcus sanguinis
pentose-phosphate shunt
lactose catabolic process via tagatose-6-phosphate
glycine betaine transport
regulation of DNA replication
cellular metabolic compound salvage
DNA unwinding involved in DNA replication
DNA repair
response to oxidative stress
cell cycle
fructose 6-phosphate metabolic process
phenylalanyl-tRNA aminoacylation
iron-sulfur cluster assembly
protein refolding
adenine salvage
cell redox homeostasis
protein import
protein repair
cytolysis
polysaccharide catabolic process
0% vs. 5%
0% vs. 50%
5% vs. 50%

**Figure S8**. **Bar graphs summarize the expression of nitrosative stress response genes across serum concentrations.** The plots show the total number of upregulated and downregulated genes, previously identified in in vitro studies, that were differentially expressed, as well as the subset exhibiting the same expression trend in response to serum. The set of nitrosative stress response genes were identified by previous *in vitro* studies (Belvin et al 2019 [46], Lewis et al 2012 [47]).

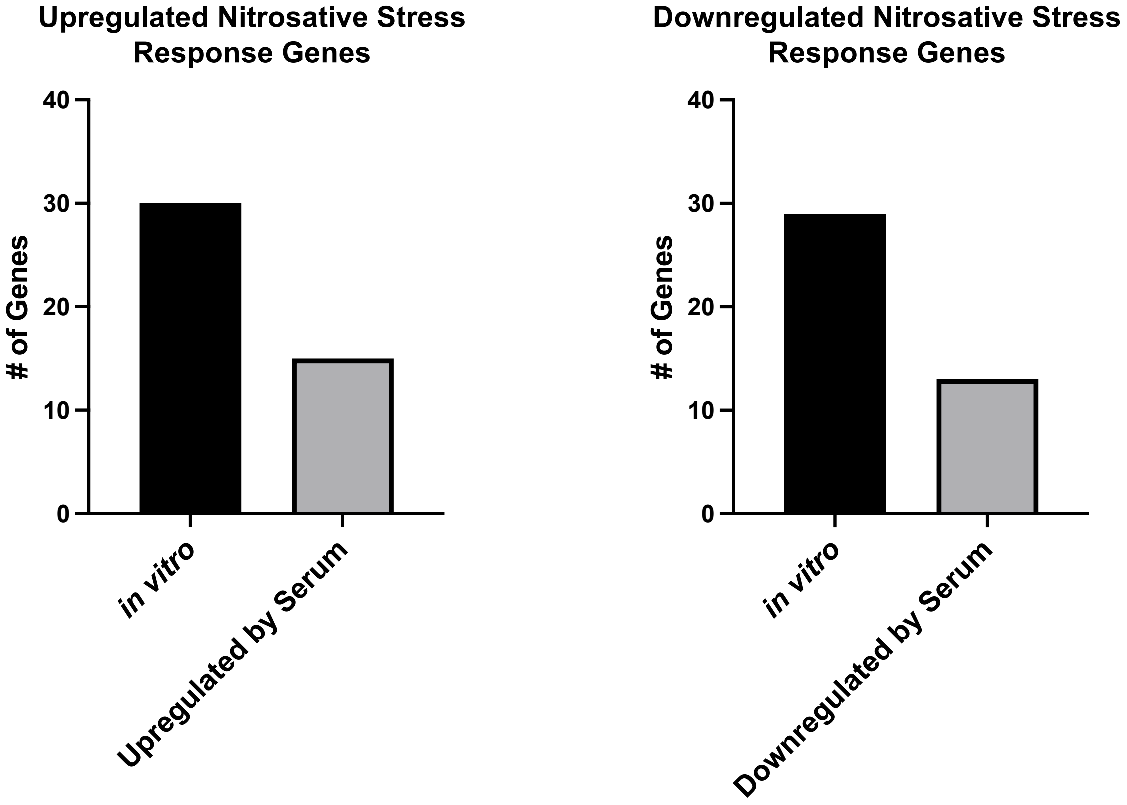
